# CD40 Signaling Restricts Retrograde Viral Spread and Provides Neuroprotection to Retinal Ganglion Cells in a Murine β-Coronavirus Model of Optic Neuritis

**DOI:** 10.64898/2026.08.18.745465

**Authors:** Niveditha E, Bishal Hazra, Souvik Karmakar, Subhajit Das Sarma, Kenneth S Shindler, Jayasri Das Sarma

## Abstract

CD40, a co-stimulatory receptor of the tumor necrosis factor receptor superfamily expressed on microglia and macrophages, is an upstream regulator of innate antiviral defense in coronavirus-induced neuroinflammation, but its specific role in the visual system remains undefined. Here, we demonstrate that CD40 signaling is essential for restricting retrograde axonal transport of the murine β-coronavirus RSA59 from the brain to the retina and for preventing chronic neurodegeneration in a model of viral optic neuritis. Wild-type (WT) and CD40−/− mice were intracranially inoculated with RSA59, and viral burden, neuroinflammation, and neurodegeneration were assessed at acute (day 5), bridging (day 7), and chronic (day 30) stages. CD40−/− mice exhibited significantly increased clinical severity and ∼30% mortality by day 12 post-infection (p.i.), compared to 100% survival in WT mice. CD40 deficiency resulted in elevated viral loads in the optic nerve and enhanced retrograde viral dissemination across all retinal layers, whereas in WT mice, the virus was largely confined to the ganglion cell layer. CD40−/− mice exhibited impaired early microglial activation and compensatory astrogliosis during the acute and bridging phases. By day 30 p.i., although viral nucleocapsid protein was undetectable by immunohistochemistry in both genotypes, CD40−/− optic nerves retained significantly higher persistent viral RNA and exhibited extensive demyelination, oligodendrocyte loss, axonal depletion, and upregulation of phagocytic markers. Critically, CD40−/− retinas showed persistent astrogliosis, accumulation of phagocytic microglia/macrophages, and a significant loss of Brn3a+ retinal ganglion cells. These findings establish CD40 as a critical molecular node governing coronavirus optic neuritis, linking early innate immune regulation to long-term neuronal survival.

**Importance:** Coronaviruses can invade the nervous system and cause lasting neurological damage, but the host immune mechanisms that limit viral spread within neural circuits are poorly understood. This study identifies CD40, a receptor on immune cells in the nervous system, as an essential defender against the retrograde spread of murine coronavirus RSA59 from the brain to the retina along nerve cell fibers. When CD40 is absent, the virus travels unchecked into the retina, initial immune responses by microglia are blunted, and a cycle of chronic inflammation ensues, ultimately destroying the retinal ganglion cells responsible for vision. These findings reveal a critical link between early immune activation and long-term neuronal protection against virus-induced damage, providing insight into how coronavirus infections lead to persistent neurological problems and highlighting CD40 as a potential therapeutic target.

## Introduction

Optic neuritis (ON) is an inflammatory demyelinating condition of the optic nerve that represents a common and frequently debilitating early manifestation of multiple sclerosis (MS) and related neuroinflammatory syndromes(1–3). Clinical ON is characterized by inflammatory infiltration of the optic nerve, progressive demyelination, and retrograde degeneration of retinal ganglion cell (RGC) axons, ultimately culminating in permanent visual impairment(4, 5). While the autoimmune model of ON, as studied extensively in the experimental autoimmune encephalomyelitis (EAE) system, has provided important mechanistic insights(6, 7), growing evidence supports a viral etiology in a subset of MS and ON cases, necessitating dedicated virus-induced experimental models to understand the full spectrum of pathogenic mechanisms(8–11).

Mouse hepatitis virus (MHV), a neurotropic murine β-coronavirus, has been extensively employed as a model for virus-induced CNS demyelination that recapitulates key histopathological features of MS, including acute meningoencephalitis, progressive white matter demyelination, and axonal degeneration(12–14). Among the recombinant isogenic MHV strains, RSA59, a spike-gene recombinant of MHV-A59, is a demyelinating (DM) strain that induces optic neuritis, marked by macrophage infiltration to the optic nerve, demyelination, and significant axonal loss following intracranial inoculation in C57BL/6 mice; whereas the non-demyelinating (NDM) isogenic control strain RSMHV2 does not(11, 15). The differential pathogenicity between these strains depends critically on the viral spike glycoprotein, which mediates both anterograde and retrograde axonal transport and is required for the induction of optic neuritis and RGC loss(11, 15, 16). Further studies using a proline-deleted fusion peptide mutant RSA59(P) established that an intact proline dyad in the spike protein fusion peptide is necessary for efficient cell-to-cell fusion, retrograde axonal translocation to RGC bodies in the retina, and subsequent neurodegeneration. Proline mutant infected mice showed significantly reduced optic nerve demyelination and no significant RGC loss compared to wild-type RSA59(PP)-infected mice(17, 18). Collectively, these foundational studies demonstrate that RSA59 reaches the retina from the brain via retrograde axonal transport along RGC axons, a process initiated at the lateral geniculate nucleus (LGN) in the thalamus, with viral antigen detectable in optic nerves as early as day 3 and in the retinal GCL by day 6 post-infection(p.i.)(11, 19).

While these studies delineated the virological determinants of retrograde transport, the host innate immune mechanisms that restrict this dissemination and preserve the structural integrity of the visual system remain incompletely characterized. Previous work has established that innate immunity, particularly the activation of microglia and macrophages, is the predominant early immune response in RSA59-infected optic nerves, with Iba1+ cells comprising the majority of infiltrating leukocytes during the acute phase, while few or no CD3+ T cells or CD19+ B cells are present at this stage(11, 20). These findings underscore the importance of the innate myeloid compartment in the early control of coronavirus-induced optic neuritis(11, 21).

A series of recent studies has delineated a hierarchically organized independent neuroimmune circuit, the Ifit2 (interferon-induced protein with tetratricopeptide repeats 2) and CD40–CD40L–CD4+ T cell axis, that is indispensable for restricting RSA59 pathogenesis in the CNS. Ifit2 (ISG54), an interferon-stimulated gene product, was identified as a critical innate regulator: Ifit2−/− mice exhibit high mortality, uncontrolled viral replication, paradoxically attenuated early microglial activation, and severe chronic demyelination and RGC loss upon RSA59 infection(22, 23). Crucially, Ifit2 has been proposed to function as a downstream effector of the CD40–CD40L co-stimulatory axis, positioning these immunological co-stimulatory signals upstream of the interferon-induced gene network(22, 23). CD40L deficiency leads to high mortality, impaired CD4+ T cell priming in cervical lymph nodes, failure of CNS lymphocyte infiltration, and severe chronic demyelination in RSA59-infected mice(24). Notably, while CD40L deficiency abrogates host protection, CD40 deficiency was predicted to produce a distinct phenotype. CD40−/− mice exhibit increased acute-phase encephalitis driven by heightened neutrophil infiltration, amplified bystander neuroinflammation, and a mechanistically distinct failure of viral control (unpublished data). This functional asymmetry within the dyad reflects that CD40, expressed on microglia, primarily modulates the magnitude and character of innate inflammatory responses, while CD40L on CD4+ T cells drives adaptive T cell priming and CNS recruitment.

Despite this emerging framework, whether CD40-dependent signaling on microglia and macrophages specifically restricts retrograde axonal transport of RSA59 into the visual system and protects retinal neurons from virus-induced neurodegeneration has never been investigated. In this study, we directly tested this hypothesis using CD40−/− mice in the established RSA59 model of viral optic neuritis. We demonstrate that CD40 deficiency allows unrestricted viral spread throughout the retina, impairs early microglial activation, paradoxically augments astrogliosis, leads to chronic viral RNA persistence, and culminates in severe demyelination, axonal loss, and RGC death. Our data identify CD40 as a non-redundant molecular component of the innate immune barrier in the visual system and provide mechanistic insight into the neuroimmune axis governing coronavirus-induced optic neuritis.

## Materials and Methods

### Animals

Age-matched five-week-old male wild-type (WT) C57BL/6 mice and CD40−/− (Jackson Laboratory, B6.129P2-Cd40tm1Kik/J) mice on the C57BL/6 background were used in all experiments. Mice were maintained under specific pathogen-free conditions in accordance with institutional guidelines. All experiments were approved by the Institutional Animal Ethics Committee (IAEC; IISERK/IAEC/AP/2024/128) and conducted in compliance with the guidelines of the Committee for the Control and Supervision of Experiments on Animals (CCSEA). Mice were monitored daily post-infection and were euthanized at prespecified time points as per the kinetics of MHV infection.

### Virus and Intracranial Inoculation

The recombinant demyelinating strain RSA59, an isogenic spike-gene recombinant of MHV-A59 expressing enhanced green fluorescent protein (EGFP), was used for all experiments(25). The recombinant virus, RSA59, was propagated in murine 17Cl-1 cells, while plaque assays and plaque purification were performed using murine L2 cells(17). WT and CD40−/− mice were inoculated intracranially with 2500 plaque-forming units (PFU) of RSA59 in a final volume of 20 μl of 0.75% Phosphate buffered saline/Bovine Serum Albumin as previously described(22, 24). Mock-infected control mice were inoculated with an equivalent volume of 0.75% PBS/BSA. The virus was injected directly into the brain via a syringe needle through the posterior skull, posterior to the orbits, near the lateral geniculate nucleus. Mice were monitored daily for clinical signs of disease.

### Clinical Scoring and Body Weight Monitoring

Mice were weighed daily and assessed for neurological signs using a standardized 0–4 clinical scoring scale as described previously(19, 22): 0, no symptoms; 0.5, numbness in the tail and hunched posture; 1, weight loss and mild ataxia; 1.5, moderate weight loss and ataxia; 2, severe weight loss and difficulty righting; 2.5, hind-limb weakness; 3, partial hind-limb paralysis; 3.5, complete hind-limb paralysis; 4, moribund or dead. Survival was recorded daily for up to day 30p.i. Data are shown for N = 5 mice per group per experiment, with at least three independent replicates.

### Tissue Harvesting and Processing

At days 5, 7 (acute and bridging phases), and 30 (chronic phase) p.i., mice were euthanized using isoflurane and transcardially perfused with 1× phosphate-buffered saline (PBS). Optic nerves and whole eyes were dissected and post-fixed in 4% PFA for 15 minutes (for optic nerve) and 14-16 hours (for eye) at room temperature, processed, and then embedded in paraffin. An automated microtome Thermo Scientific™ HM 355S was used to generate five-micron-thick longitudinal optic nerve sections, and retina sections within cross-sections of whole eyes were sectioned for subsequent histological and immunohistochemical analyses.

### Histopathological Analysis

Paraffin sections of optic nerves and retinas were stained with hematoxylin and eosin (H&E) to assess inflammatory infiltration and overall tissue architecture. Myelin integrity in optic nerve sections at day 30p.i. was evaluated using Luxol Fast Blue (LFB) staining, and demyelinated areas were quantified using Fiji/ImageJ (ImageJwin64) as previously described(18, 26). All slides were coded and evaluated in a blinded fashion by at least two independent investigators.

### Immunohistochemistry

Paraffin sections were deparaffinized, rehydrated, and subjected to Heat-induced epitope retrieval (HIER) with Antigen Unmasking Solution (Citrate-Based; Vector Laboratories, Cat. No. H-3300). Sections were blocked with appropriate serum and incubated overnight at 4°C with primary antibodies. The following primary antibodies were used: anti-nucleocapsid (N) protein of RSA59 (1:50; Kind gift from Julian Leibowitz, Texas A&M University) for viral antigen detection; anti-Iba1 (FUJIFILM Wako Chemicals, catalog no. 019-19741, 1:500) for microglia/macrophage labeling; anti-GFAP (Sigma-Aldrich, G3893) for astrocyte detection. Secondary antibody detection was performed using the avidin–biotin–immunoperoxidase technique (Vector Laboratories) with 3,3′-diaminobenzidine (DAB) as the chromogenic substrate for brightfield immunohistochemistry(27). Slides were counterstained with hematoxylin where appropriate. Images were captured using the Invitrogen EVOS XL Core microscope. Further, they were quantified using Fiji/ImageJ (ImageJwin64) as previously described(17, 28).

### Immunofluorescence

Paraffin tissue sections were deparaffinized, rehydrated, antigen-retrieved, incubated with 0.1 M glycine in PBS to reduce non-specific cross-linking, and then treated with 1mg/ml NaBH4 to reduce autofluorescence. The slides were permeabilized with 0.2% Triton X-100 in PBS by shaking incubation at room temperature (RT) for 15 min, followed by blocking with 1% BSA prepared in 0.2% Triton X-100 in 1× PBS at 37°C for 1 h. Subsequently, the sections were incubated with primary antibodies diluted in blocking solution under shaking conditions at 4°C for 16h(23, 29). Individual sections from each sample slide were independently stained using the following antibody combinations: anti-MBP (homemade serum antibody, 1:1) and anti-PLP (homemade serum antibody, 1:1) for mature oligodendrocytes/myelin assessment; anti-neurofilament medium chain (Sigma-Aldrich, catalog no. N5389, 1:100) for axonal integrity; anti-Iba1 (FUJIFILM Wako Chemicals, catalog no. 019-19741, 1:500) for microglia/macrophage labeling; and anti-Brn3a (Synaptic Systems, catalog no. 411003, 1:500) for RGC quantification. The slides were subsequently washed and incubated for 1 h at 37°C with Alexa Fluor 568-conjugated donkey anti-rabbit (Invitrogen, A10042) and Alexa Fluor 488-conjugated goat anti-mouse (Invitrogen, A11001) secondary antibodies, each diluted in blocking solution (1:750). Nuclei were counterstained with DAPI. Images were captured using an Olympus IX-81 microscope operating at 40x magnification. Quantification of Brn3a immunoreactivity was performed by calculating the mean fluorescence intensity (MFI) using ImageJ (NIH, USA).

### Quantitative RT-PCR

For viral RNA quantification, optic nerves (10 optic nerves from five mice pooled per group) and retinas (10 eyes from five mice pooled per group) were harvested at days 5, 7, and 30 p.i. Total RNA was extracted using TRIzol reagent. Total RNA concentration was quantified using a NanoDrop™ 2000 Spectrophotometer. cDNA was synthesized from 1 µg of total RNA using the High-Capacity cDNA Reverse Transcription Kit (Applied Biosystems, Cat. No. 4368814). Quantitative real-time PCR (qRT-PCR) was performed using iTaq™ Universal SYBR® Green Supermix on a Bio-Rad C1000 Touch Thermal Cycler under the following cycling conditions: initial denaturation at 95°C for 10 min, followed by 40 cycles of denaturation at 95°C for 15 s and annealing/extension at 60°C for 1 min. Melt curve analysis was subsequently performed at 60°C for 1 min. All reactions were run in triplicate in 96-well plates with an automatically assigned baseline and manually adjusted threshold cycle (Ct) values and analyzed using the 2−ΔΔCT method. Dissociation curve analysis confirmed the amplification specificity and presence of a single PCR product for each primer pair in every sample. Primer sequences are listed in Table 1.

**Table 1:**
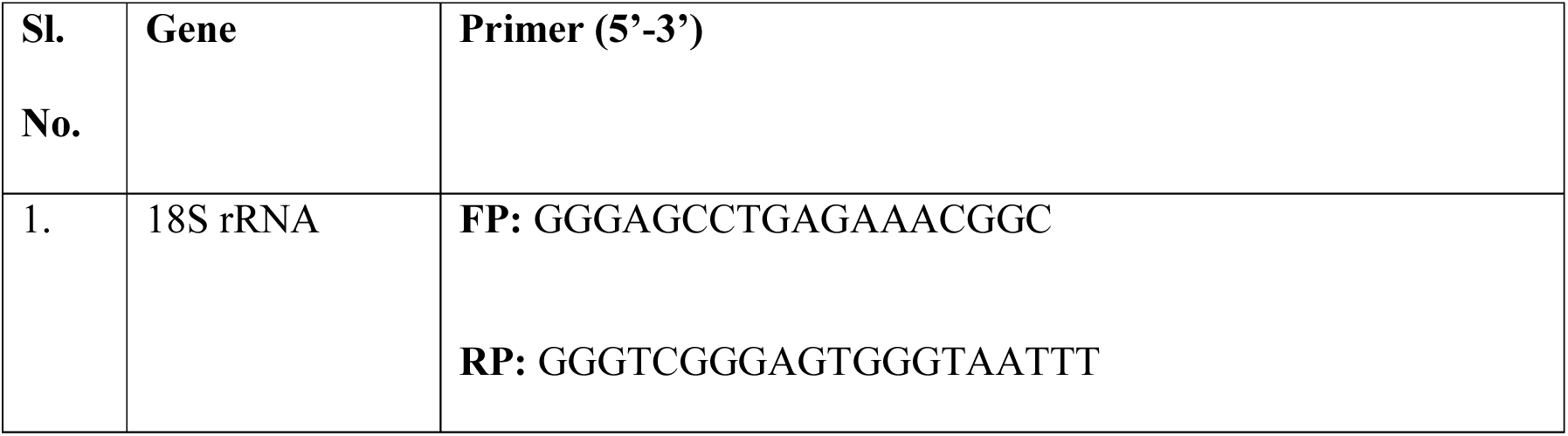
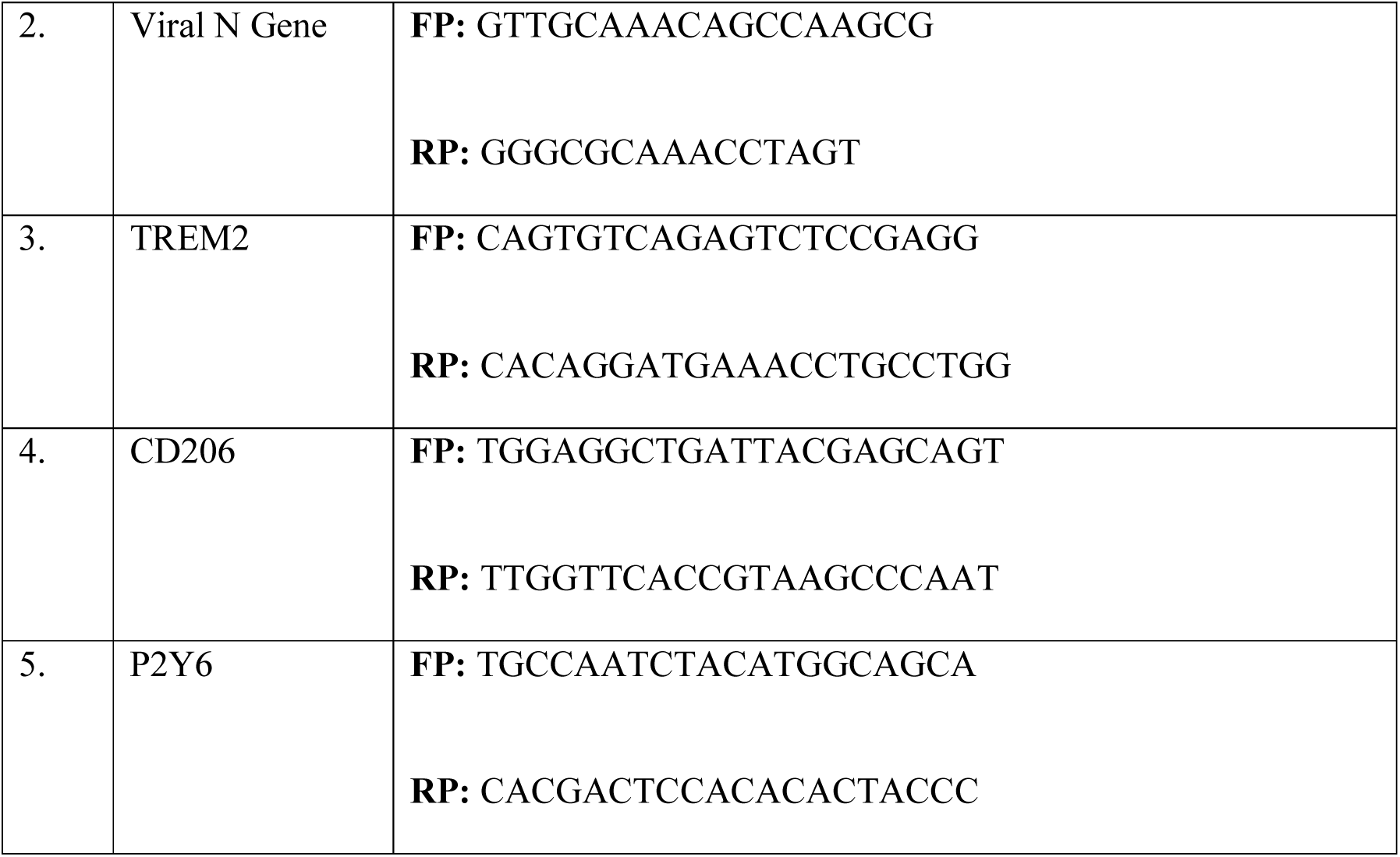
List of Primers used.

### Statistical Analysis

Comparisons between WT and CD40−/− groups at individual time points were analyzed using an unpaired Student’s t-test. Survival curves were compared using the log-rank (Mantel–Cox) test. All data are presented as mean ± SEM. Statistical significance was set at p < 0.05. Analyses were performed using GraphPad Prism software. Statistical notations used: *p<0.05, **p<0.01, ***p<0.001, ****p<0.0001.

## Results

### CD40 Deficiency Increases Susceptibility to RSA59 Infection

To determine whether CD40 deficiency alone contributes to baseline pathological changes in the optic nerve and retina, wild-type (WT) and CD40−/− mice were mock-infected via intracranial inoculation with uninfected cell lysate (PBS supplemented with 0.075% BSA). No pathological changes in optic nerve or retina were observed, confirming that CD40 deficiency does not cause baseline structural abnormalities in the visual system (Fig. 1). Hence, to determine the role of CD40 in the host immune response to neurotropic coronavirus infection, WT and CD40−/− mice were intracranially inoculated with 2500 PFU of RSA59 and monitored daily. WT mice maintained 100% survival throughout the observation period, exhibiting only mild, transient clinical signs (scores 0.5–1) and minimal weight loss. In contrast, CD40−/− mice developed significantly more severe clinical disease, with a progressive and marked decrease in body weight beginning at day 7p.i. and substantially elevated clinical scores compared to WT mice at all post-infection time points (Fig. 2A, B). Survival analysis of CD40-/- mice demonstrated a significantly lower survival rate of 60% by day 10 p.i, compared to 0% mortality in WT controls (Fig. 2C). Mock-infected WT and CD40−/− mice displayed no major phenotypic differences, aside from slightly smaller body sizes in the CD40-/- pups. These data establish that CD40 is an important determinant of clinical resistance to RSA59 infection.

**Figure 1:**
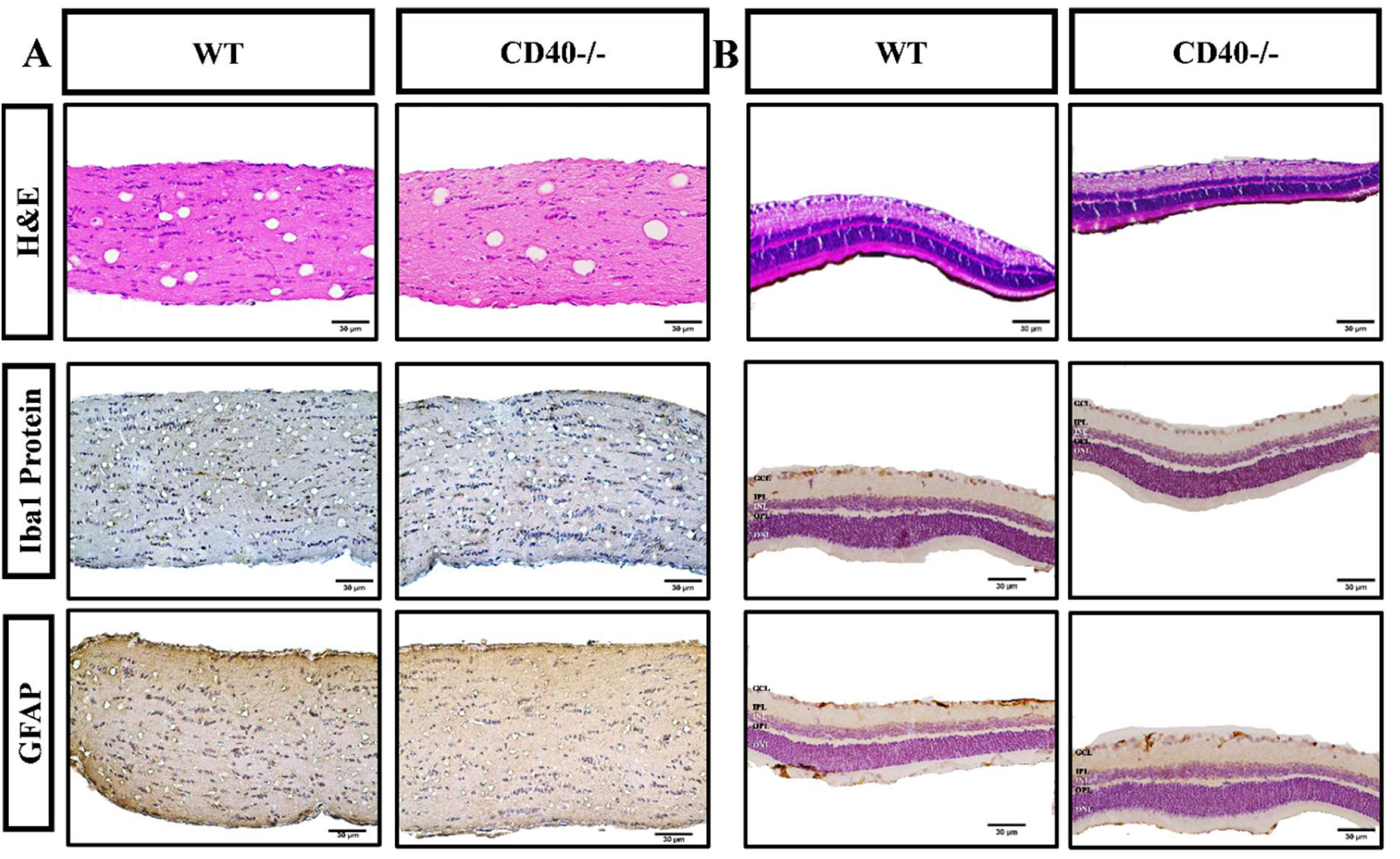
Absence of CD40 does not induce any detectable pathology in mock-infected mice. Optic nerve and retina cross-sections from 0.75% PBS-BSA-infected (Mock), WT, and CD40-/- mice were collected and stained with Hematoxylin & Eosin, and immunohistochemically with anti-Iba1 and anti-GFAP. No observable differences were found between the WT and CD40-/- mice samples of optic nerve (A) and retina (B), indicating that CD40 deficiency does not cause any pathological changes in mock-infected samples. N=3 per group for histopathological experiments.

**Figure 2:**
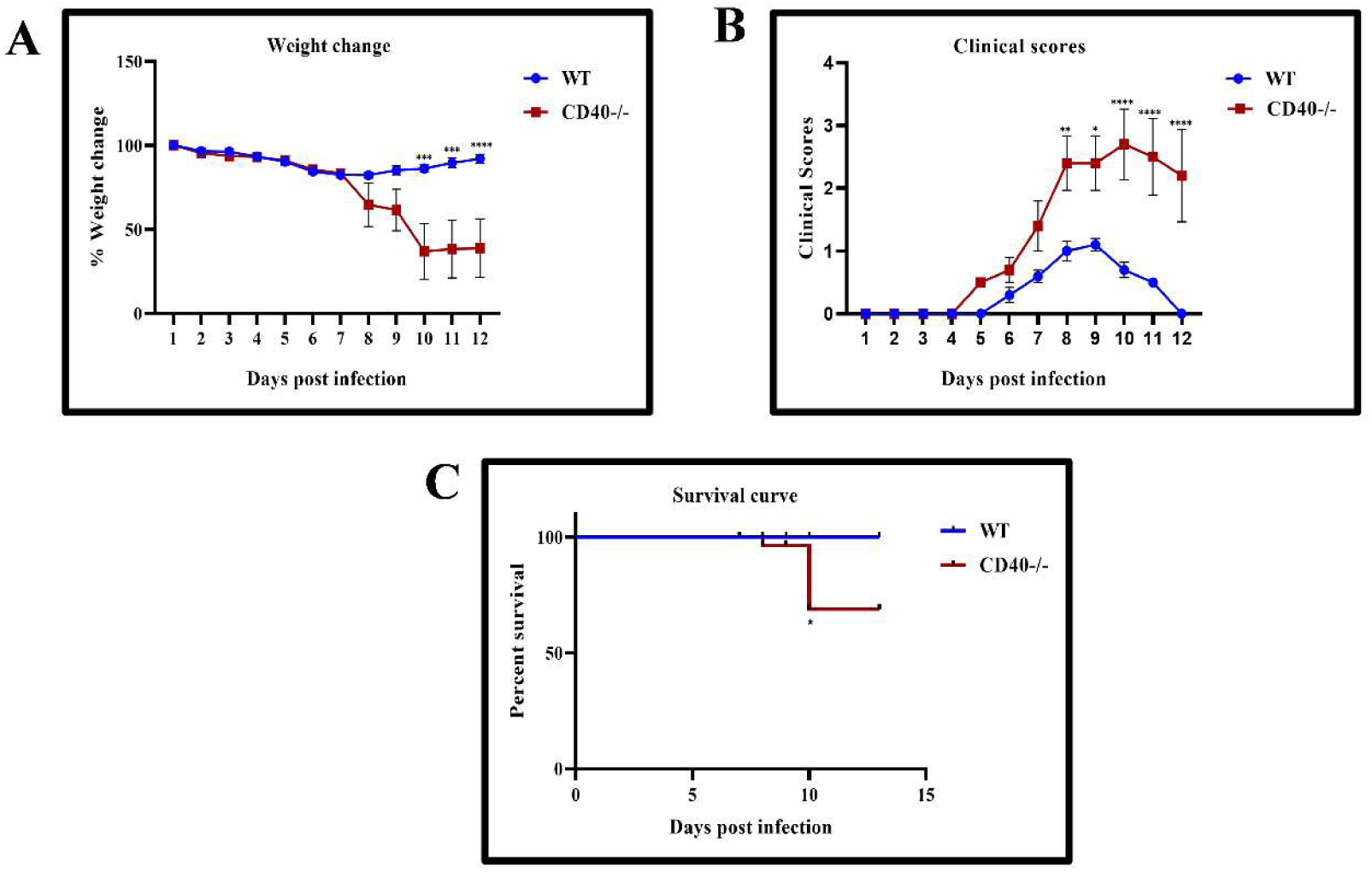
CD40 Deficiency increases susceptibility to RSA59 Infection. 5-week-old WT (N = 38) and CD40^-/-^ mice (N = 42) were infected with RSA59 (2500 PFUs) and monitored daily for (A) weight change, (B) clinical scores, and (C) survivability. Clinical scores were assigned on a 0–4 scale, as described in the Materials and Methods section. *Asterisk represents statistical significance calculated using an unpaired Student’s t-test; p<0.05 was considered significant. Statistical significance for the survival curve was determined by the Log-rank (Mantel-Cox) test. *p<0.05, **p<0.01, ***p<0.001, ****p<0.0001.

### CD40 Deficiency Results in Increased Viral Persistence in the Optic Nerve during the Acute and Bridging Phases

To assess whether CD40 contributes to the restriction of RSA59 spread within the visual axis, optic nerve sections from WT and CD40−/− mice were immunostained for viral nucleocapsid (N) protein at days 5 and 7 p.i. (Fig. 3). CD40−/− mice exhibited significantly higher areas of N-protein-positive staining throughout the optic nerve at both day 5p.i. and day 7p.i. compared to WT mice (Fig. 3A–C, E–G). This enhanced viral burden was corroborated by qRT-PCR analysis of pooled optic nerve samples, which revealed elevated viral N-gene transcript levels in CD40−/− mice at both acute and bridging time points (Fig. 3D, H). These findings demonstrate that CD40-dependent innate immune signaling is required for the early restriction of RSA59 dissemination within the optic nerve.

**Figure 3:**
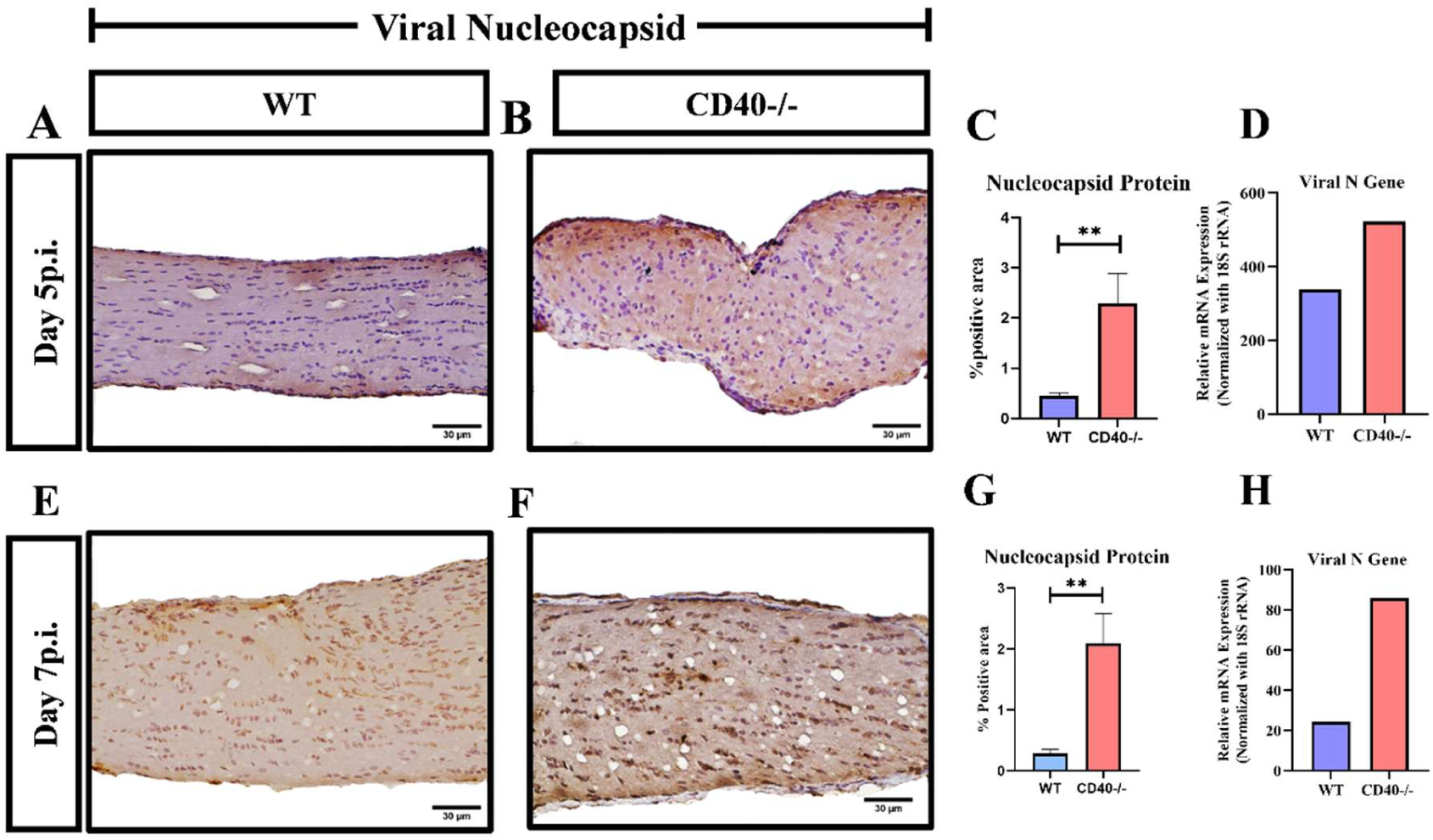
Absence of CD40 signaling results in elevated viral persistence throughout acute and bridging disease phases. 5μm thin paraffin sections were prepared from optic nerves of 5-week-old WT and CD40-/- infected mice at days 5 and 7 p.i, and immunohistochemically stained for viral antigen using viral anti-N (Nucleocapsid) antibody. CD40-/- mice showed increased viral spread at both day 5 p.i. and day 7 p.i. (panel B and F, respectively) as compared with the WT mice (panel A and E, respectively). Quantification of Viral N protein expression is shown in Panels C and G. Increased Viral N gene expression was also observed in CD40-/- at day 5 p.i. (Panel D) and day 7 p.i. (Panel H) mice optic nerve (10 optic nerves from 5 mice pooled together in each group). The WT and CD40-/-mice’s optic nerve regions were magnified at 40X, and the scale bar for all sections is 30 μm. Data are represented as mean ± SEM (N=3 per group, for immunohistochemistry). Statistical significance was determined using an unpaired T test; **p < 0.01.

### CD40 Deficiency Enhances Retrograde Axonal Transport of RSA59 to the Retina and Promotes Pan-Retinal Viral Spread

Prior studies established that RSA59 reaches the retina from the brain via retrograde axonal transport along RGC axons, with viral antigen initially detectable in the GCL and subsequently spreading only to a limited extent to deeper retinal layers (11, 18, 19). To determine whether CD40 restricts this process, retinal cross-sections from WT and CD40−/− mice were immunostained for N protein at days 5 and 7 p.i. (Fig. 4). In WT mice, as expected, viral antigen was largely restricted to the GCL at day 5 p.i. and was substantially reduced by day 7p.i. In striking contrast, CD40−/− mice exhibited dramatically enhanced retrograde transport, with viral antigen spreading across all retinal layers, including the inner nuclear layer (INL), outer plexiform layer (OPL), and outer nuclear layer (ONL) at both day 5 and day 7 p.i. (Fig. 5A–H). Quantification confirmed significantly higher viral N-protein staining area in CD40−/− retinas compared to WT at both time points (Fig. 5C, G), and this was corroborated by elevated viral N-gene expression by qRT-PCR (Fig. 5D, H). These data establish that CD40 is a critical molecular barrier that impedes the retrograde axonal dissemination of RSA59 from the optic nerve into the deep retinal parenchyma.

**Figure 4:**
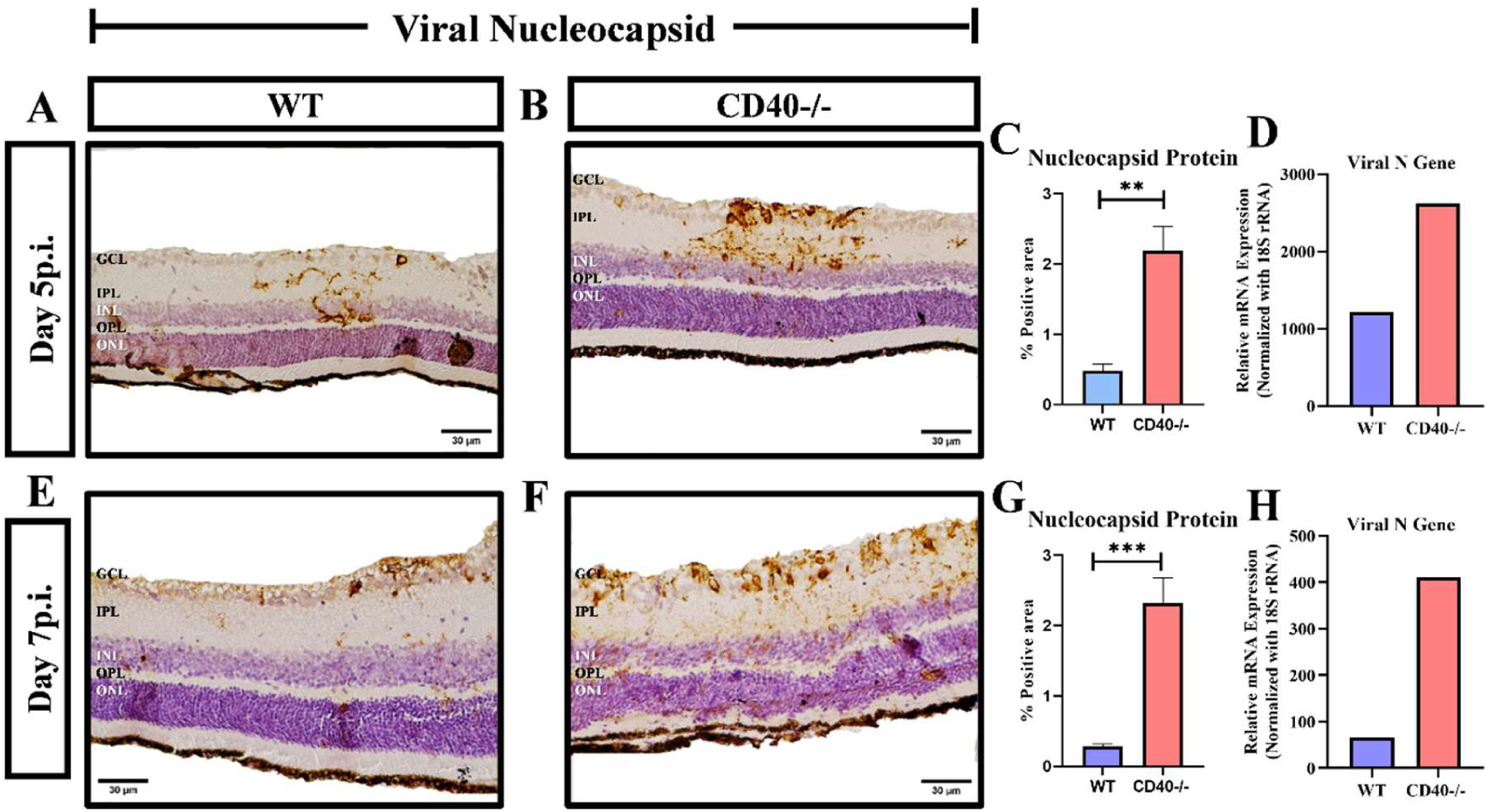
CD40 deficiency enhanced RSA59 retrograde axonal transport to the retina, promoting spread across retinal layers. 5μm thin paraffin sections were prepared from the retina of 5-week-old WT and CD40-/- infected mice at day 5 (Panel A, B) and day 7p.i (Panel E, F), and immunohistochemically stained for viral antigen using anti-N (Nucleocapsid) antibody. Quantitative analysis of viral N protein expression is presented in Panels C and G, confirming significantly higher viral burden in CD40⁻/⁻ retina. Moreover, increased viral N gene expression was also observed in the CD40⁻/⁻ mouse retina at day 5 p.i. (Panel D) and day 7p.i. (Panel H) (10 eyes from 5 mice pooled together in each group). Representative images were magnified at 40×, and the scale bar = 30 µm. Data are represented as mean ± SEM (N=3-4 per group). Statistical significance was determined using an unpaired T test; **p < 0.01 and ***p < 0.001.

**Figure 5:**
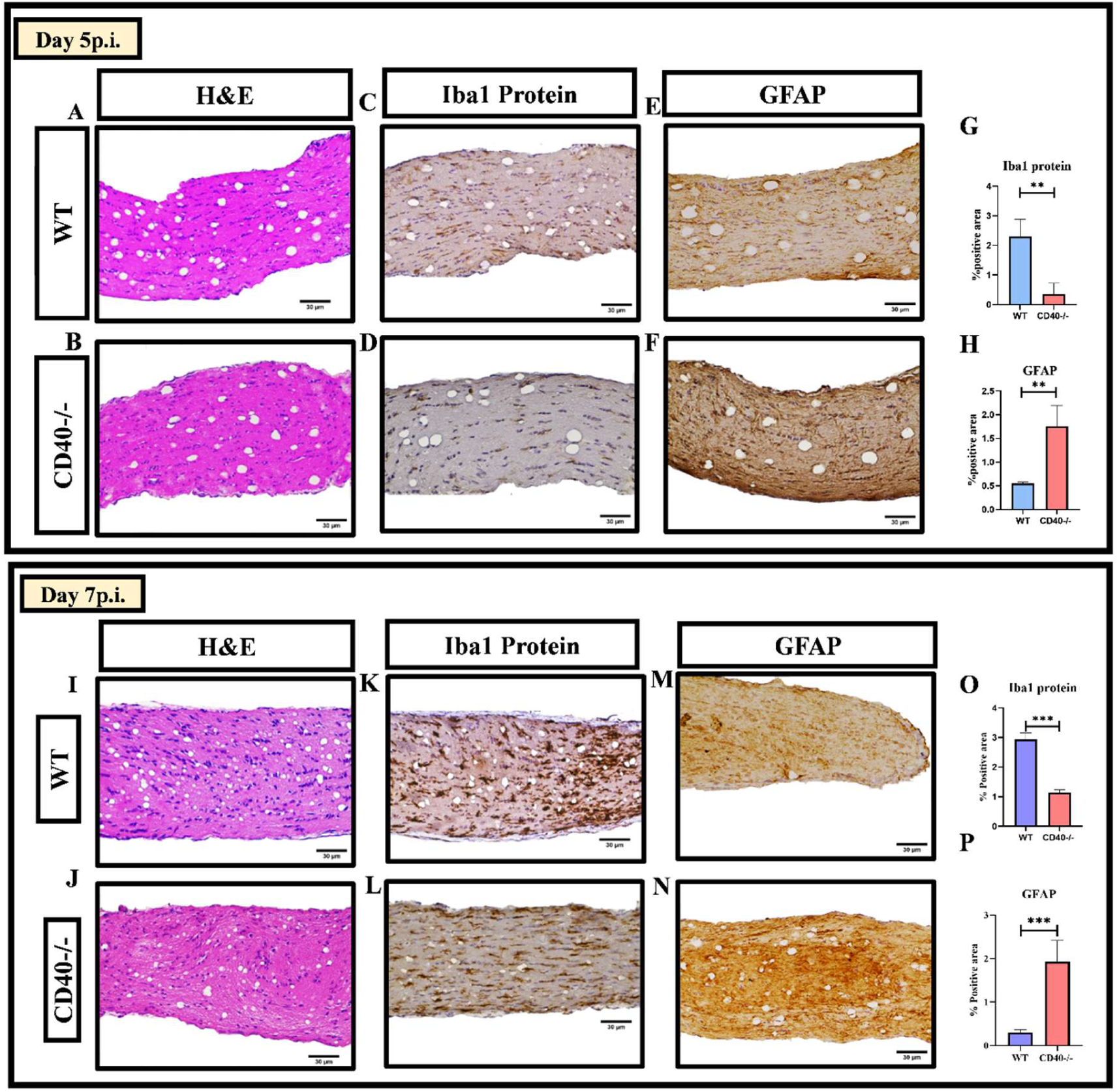
Differential neuroinflammatory responses in WT and CD40⁻/⁻ mouse optic nerves during acute & acute-chronic bridging stages. 5μm thin paraffin sections were prepared from the optic nerve of 5-week-old WT and CD40-/- infected mice at day 5 and day 7 p.i., and were stained with Hematoxylin & Eosin, and also immunohistochemically stained for Iba1 and GFAP. At day 5 p.i. and day 7p.i. H&E staining indicated increased inflammatory infiltration in WT (A, I) compared with CD40-/- mice (B, J). Iba1 staining showed decreased and impaired microglia/macrophage activation in CD40⁻/⁻ mice (D, L), compared to WT (C, K), and GFAP staining indicated enhanced astrocyte activation in CD40-/- (F, N) compared to WT mice (E, M). (G, O) and (H, P) represents the quantitative analysis of Iba1 and GFAP staining, respectively. Scale bars = 30µm. Data are represented as mean ± SEM (N=3 per group). Statistical significance was determined using an unpaired T test; **p < 0.01, and ***p < 0.001.

### CD40 Deficiency Results in Impaired Microglia/Macrophage Activation and Paradoxically Elevated Astrogliosis in the Optic Nerve and Retina during Acute and Bridging Phases

To determine the mechanistic basis for uncontrolled viral spread in CD40−/− mice, we assessed the early neuroinflammatory response in the optic nerve (Fig. 5) and retina (Fig. 6). H&E staining of optic nerve sections at days 5 and 7 p.i. revealed increased inflammatory infiltration in WT mice compared to CD40−/− mice, despite the latter harboring higher viral loads (Fig. 4A, B, I, J). Microglia represent a major component of the resident glial population in the retina. As development progresses, they establish characteristic spatial distributions within the inner plexiform layer (IPL) and the outer plexiform layer (OPL)(30). Immunostaining for Iba1 demonstrated significantly reduced and morphologically attenuated microglial/macrophage activation in CD40−/− optic nerves compared to WT at both acute time points (Fig. 4C–G, K–O; quantification in 4G, 4O). Strikingly, in parallel with impaired microglial activation, GFAP immunostaining revealed markedly enhanced astrocyte activation in CD40−/− optic nerves compared to WT at days 5 and 7p.i. (Fig. 4F, N; quantification in 4H, 4P). This differential glial response, impaired microglial/macrophage activation concurrent with pronounced astrogliosis, represents a hallmark of the innate immune failure induced by CD40 deficiency. Identical biphasic glial responses were observed in the retina: CD40−/− retinas showed significantly reduced Iba1+ cells with a failure of microglial mobilization to sites of viral entry, while GFAP expression was markedly elevated compared to WT at both day 5 and day 7 p.i. (Fig. 6). These data suggest that CD40 is required for the timely microglial activation that constitutes the first line of defense against retrograde viral spread in the visual system, and that astrogliosis serves as a compensatory but insufficient response when CD40-mediated microglial signaling is absent.

**Figure 6:**
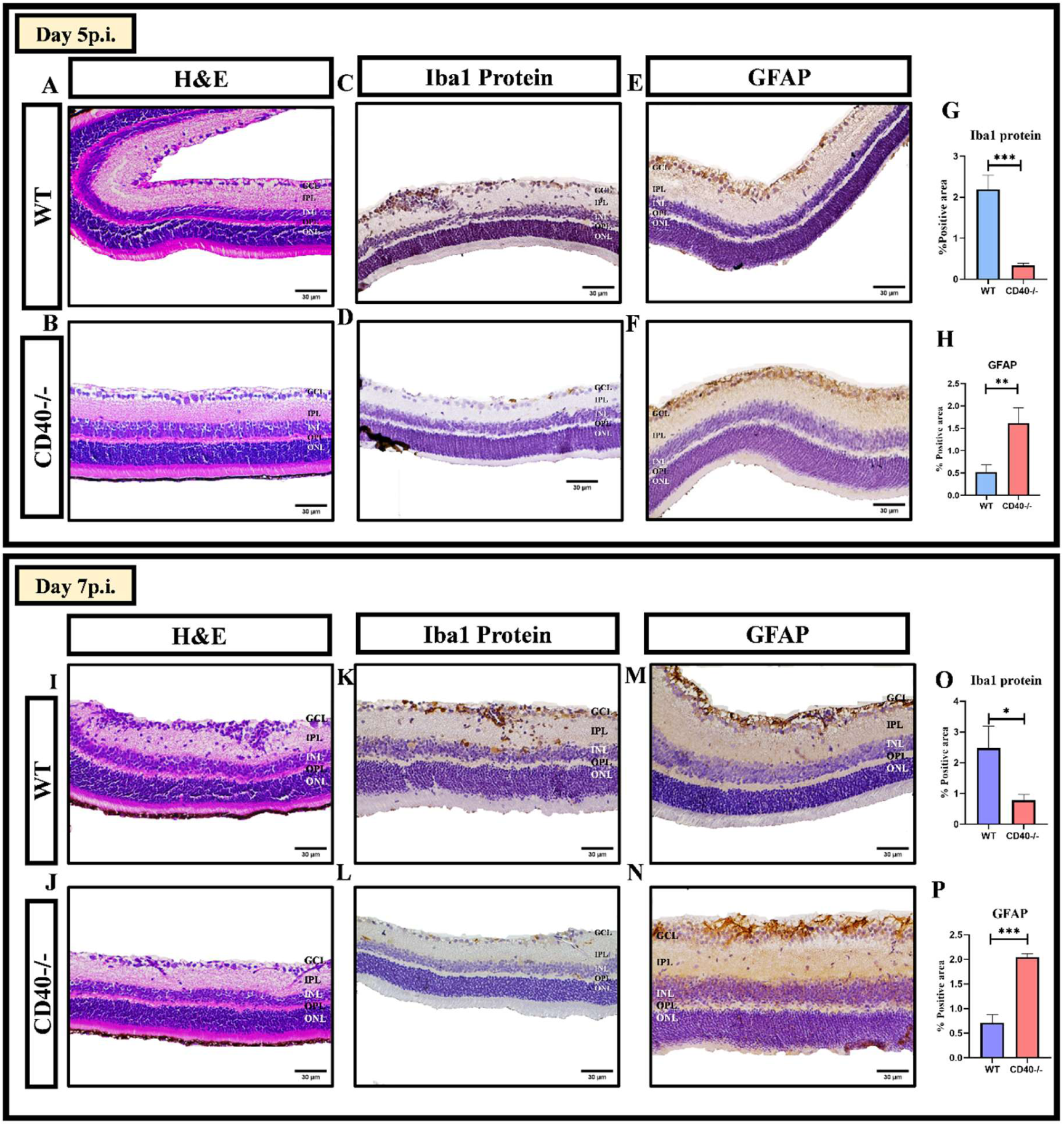
Differential neuroinflammatory responses in WT and CD40⁻/⁻ mouse retinas during acute & acute-chronic bridging stages. 5μm thin paraffin sections were prepared from the retina of 5-week-old WT and CD40-/- infected mice at day 5 and day 7 p.i., and were stained with Hematoxylin & Eosin, and also immunohistochemically stained for Iba1 and GFAP. At day 5p.i. and day 7p.i. H&E staining indicated increased inflammatory infiltration in WT (A, I) as compared to CD40-/- mice (B, J). Iba1 staining showed decreased and impaired microglia/macrophage activation in CD40⁻/⁻ mice (D, L), as compared to WT (C, K), and GFAP staining indicated enhanced astrocyte activation in CD40-/- (F, N) compared to WT mice (E, M). (G, O) and (H, P) represents the quantitative analysis of Iba1 and GFAP staining, respectively. Scale bars = 30µm. Data are represented as mean ± SEM (N=3 per group). Statistical significance was determined using an unpaired T test; *p<0.05, **p<0.01, ***p<0.001.

### CD40 Deficiency Leads to Persistent Viral RNA in the Optic Nerve during the Chronic Stage Despite Absence of Detectable Nucleocapsid Protein

At day 30p.i., immunohistochemical staining for N protein revealed no detectable viral antigen in either WT or CD40−/− optic nerve sections, consistent with the resolution of productive infection (Fig. 7A, B). However, qRT-PCR analysis demonstrated that CD40−/− mice harbored higher levels of viral N-gene RNA in the optic nerve compared to WT mice at this chronic time point (Fig. 7C). This persistent sub-threshold viral RNA in the absence of detectable antigen is analogous to findings reported previously in the CD40L−/− RSA59 model and indicates that CD40 deficiency prevents complete viral clearance, leaving a reservoir of viral genomic material that likely drives persistent inflammatory signaling and chronic neurodegeneration.

**Figure 7:**
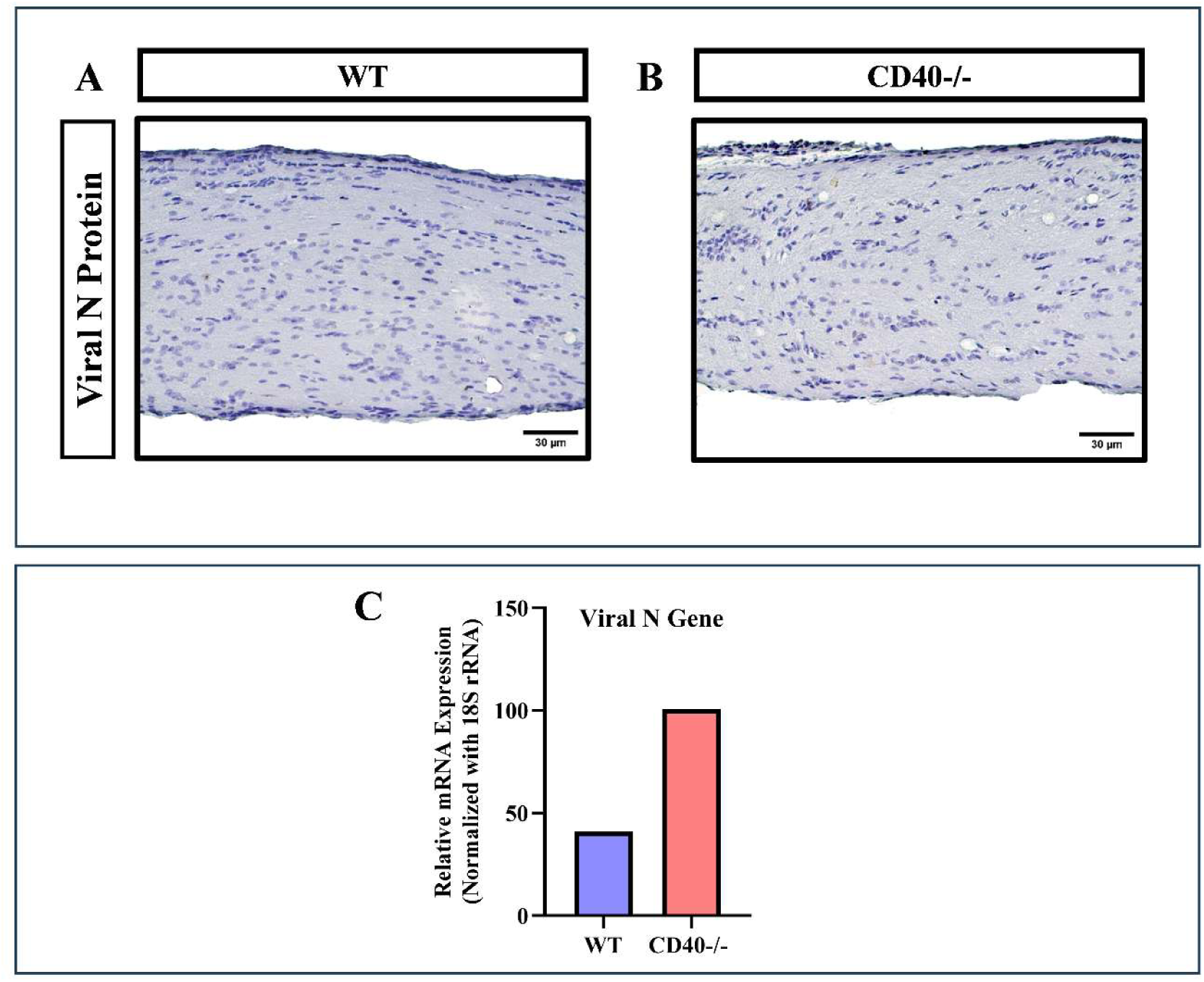
CD40 deficiency causes significant persistent viral RNA in the optic nerve without detectable nucleocapsid protein spread. During the Chronic Stage, Day 30p.i., 5 μm thin paraffin sections of RSA59-infected WT and CD40-/- mouse optic nerves were checked for the presence of viral antigen by immunohistochemical staining with Anti-N (Nucleocapsid) and showed no viral antigen in both WT (A) and CD40-/- (B). However, higher viral N gene expression was observed by qRT-PCR in CD40-/- infected mice compared with WT (C) (10 optic nerves pooled from 5 mice in each group). Scale bars = 30µm.

### CD40 Deficiency Drives Chronic Demyelination, Phagocytic Microglial Phenotypic Shift, and Sustained Astrogliosis in the Optic Nerve

Luxol Fast Blue staining of day-30p.i. optic nerve sections from CD40−/− mice revealed extensive demyelinated lesions, in marked contrast to the largely intact myelin architecture observed in WT optic nerves (Fig. 8A–C). This was accompanied by a dramatic shift in the microglial phenotype: whereas WT optic nerves showed resting or moderately activated Iba1+ microglia, CD40−/− optic nerves exhibited large numbers of activated, amoeboid phagocytic microglia/macrophages (Fig. 8D-F). In parallel, GFAP expression was significantly elevated in CD40−/− optic nerves at day 30p.i., indicating persistent, unresolved reactive astrogliosis (Fig. 8F-I). To characterize microglial phagocytic state more precisely, qRT-PCR was performed on pooled optic nerve samples using established phagocytic markers. CD40−/− mice showed upregulation of P2Y6, TREM2, and CD206 compared to WT controls (Fig. 8J–L). The collective expression of these markers indicates that microglia in CD40−/− optic nerves are engaged in aberrant, persistent phagocytic activity more consistent with myelin stripping, clearance of damaged cell debris, and tissue damage than with productive viral clearance, mirroring the phagocytic microglial phenotype observed in chronically demyelinating white matter in the RSA59 spinal cord model(14, 23, 24, 29, 31).

**Figure 8:**
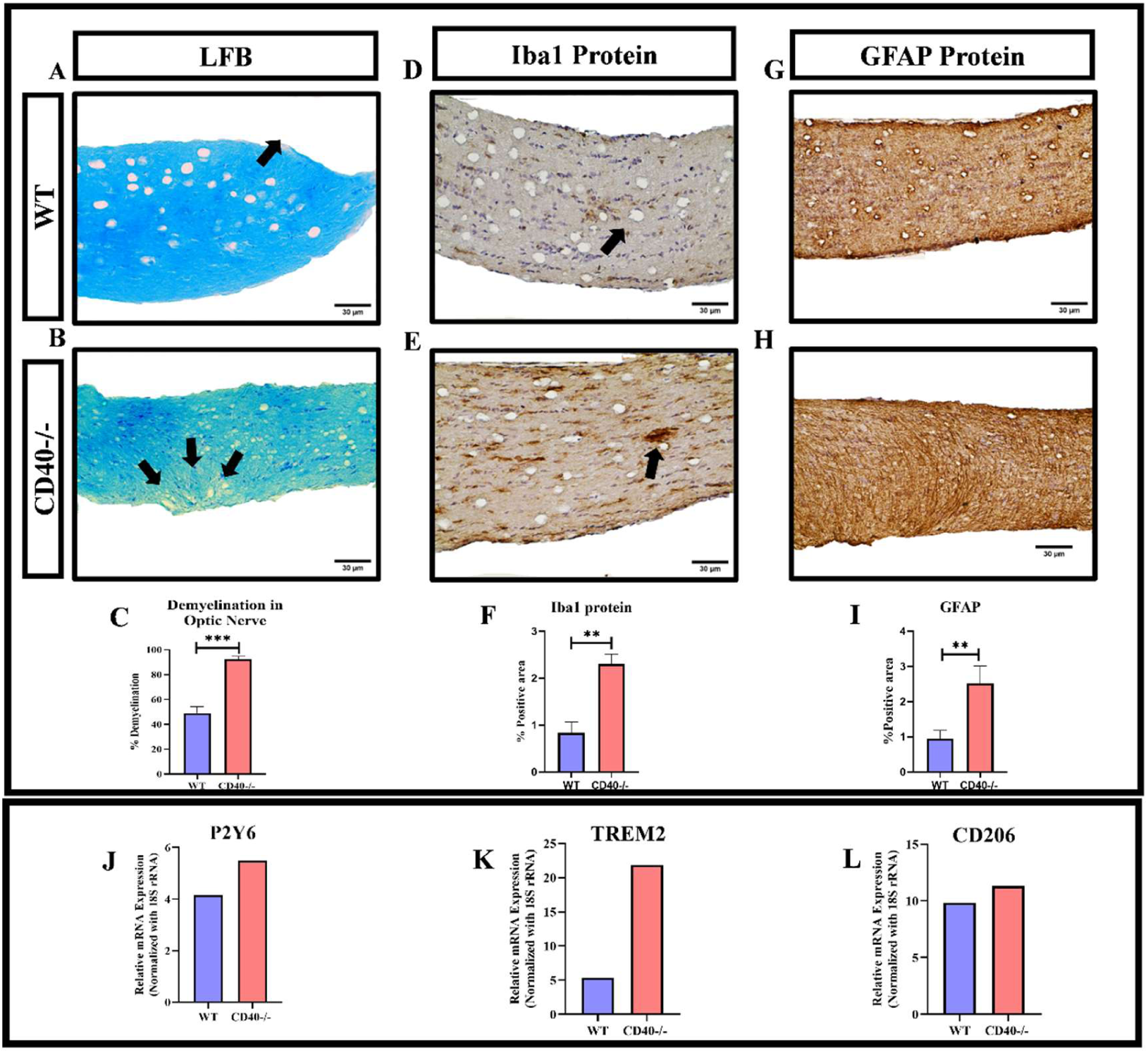
CD40 deficiency drives enhanced demyelination, promotes a phagocytic Microglia/Macrophage phenotypic shift, and sustains Astrocyte activation. At day 30p.i. 5μm-thick paraffin sections of RSA59-infected WT and CD40-/- mouse optic nerves were checked for demyelinated lesions using Luxol fast blue staining. Representative images show increased demyelinated areas (indicated by arrows) in CD40⁻/⁻ optic nerve sections (B) compared with WT (A). Microglia/macrophage activation was assessed by immunohistochemical staining with anti-Iba1. An increased presence of phagocytic microglia/macrophages in the CD40-/- optic nerve (E) compared to WT (D) was observed. A significant increase in GFAP expression was also observed in CD40⁻/⁻ mice (H) compared to WT (F), indicating persistent astrocytic activation. Quantification of LFB, Iba1, and GFAP protein expression is graphically represented in Panels C, F, and I, respectively. An upregulation of phagocytic markers, including P2Y6 (J), TREM2 (K), and CD206 (L), was observed in the optic nerves of CD40⁻/⁻ infected mice. This was determined by pooling samples (10 optic nerves from 5 mice pooled together in each group) from both groups, analyzing gene expression by qRT-PCR, and comparing WT and CD40⁻/⁻ groups. The optic nerve regions of WT and CD40-/- mice were magnified at 40X, and the scale bar for all sections is 30 μm. Statistical significance was determined using an unpaired T test; **p<0.01, ***p<0.001.

### CD40 Deficiency Leads to Loss of Mature Oligodendrocytes/Myelin-Specific Protein Expression in the Optic Nerve

To further characterize the extent of chronic white matter injury, longitudinal sections of day- 30 optic nerves from WT and CD40−/− mice were co-stained for Iba1 (microglia/macrophages) and the mature oligodendrocyte markers MBP (myelin basic protein) and PLP (proteolipid protein) (Fig. 9). WT optic nerves demonstrated preserved MBP and PLP immunoreactivity consistent with intact myelination. In contrast, CD40−/− optic nerves showed severe loss of both MBP and PLP staining in regions of active microglial/macrophage accumulation, confirming that the chronic inflammatory environment in these mice results in extensive oligodendrocyte/myelin loss and demyelination (Fig. 9A, B). These data collectively demonstrate that CD40 is required for the suppression of chronic microglia-driven demyelination, preservation of oligodendrocyte viability, and maintenance of the myelin sheath.

**Figure 9:**
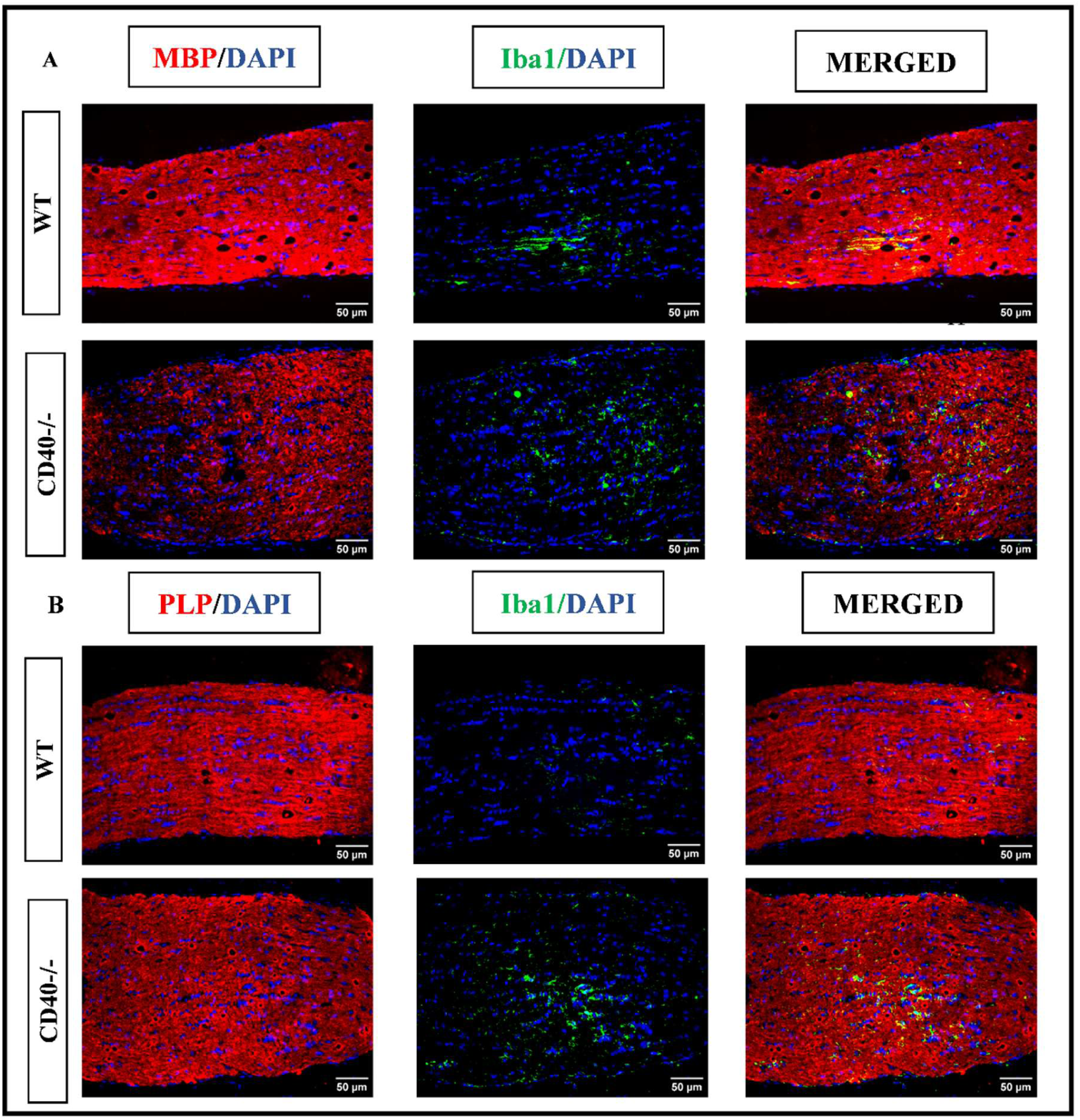
CD40 deficiency leads to loss of mature oligodendrocytes in the optic nerve. 5μm longitudinal paraffin sections were prepared from optic nerves of 5-week-old WT and CD40^-/-^ day 30 post RSA59-infected mice. Sections were stained for Iba1 (green), MBP and PLP (red), and nuclei using DAPI (blue). Representative images show areas of demyelination, indicated by loss of MBP (panel A) and PLP (panel B), together with Iba1-stained activated microglia/macrophages. Merged images show areas of overlapping staining for Iba1 and MBP, or for Iba1 and PLP. The scale bar for optic nerve sections is 50 µm. Showing one representative image from one of the three mice in one of the three independent experiments.

### CD40 Deficiency Causes Severe Axonal Loss in the Optic Nerve at the Chronic Stage

Axonal integrity was assessed on day 30p.i. optic nerves by co-labeling for Iba1 and neurofilament medium polypeptide (NFM). WT optic nerves displayed robust NFM staining, indicating intact axonal architecture (Fig. 10A). CD40−/− mice, however, showed dramatically reduced NFM immunoreactivity with large areas of axonal depletion, indicative of severe axonal degeneration (Fig. 10B). Areas of maximal axonal loss colocalized with regions of heavy Iba1+ microglial/macrophage accumulation, suggesting that the phagocytic microglial activity in CD40−/− optic nerves actively contributes to ongoing axonopathy. These findings are consistent with the concept that both direct viral cytopathic effects and bystander microglial damage converge to produce irreversible axonal degeneration in the context of chronic, unresolved neuroinflammation.

**Figure 10:**
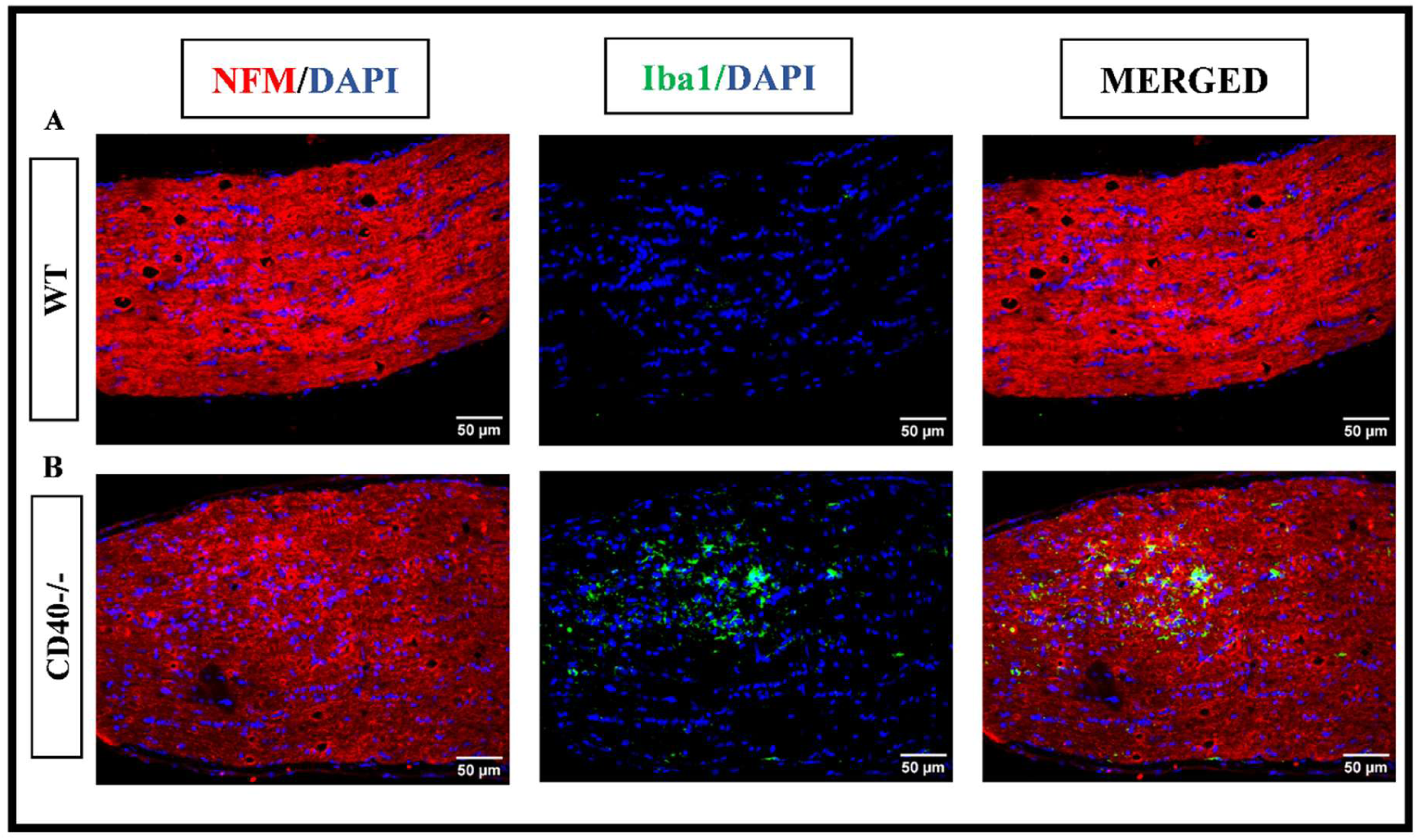
CD40 deficiency causes severe optic nerve axonal loss, evidenced by reduced neurofilament expression at the chronic stage. 5μm longitudinal paraffin sections were prepared from optic nerves of day 30 post RSA59-infected WT and CD40^-/-^ mice. Sections were stained for Iba1 (green), NFM (red), and nuclei using DAPI (blue). Representative images show increased areas of axonal loss in CD40-/- optic nerves (B), indicated by loss of neurofilament (NFM), compared with WT (A), as well as Iba1 staining for activated microglia/macrophages. Merged images show areas of overlapping staining with Iba1 and NFM, respectively. The scale bar for optic nerve sections is 50 µm. Showing one representative image from one of the three mice in one of the three independent experiments.

### CD40 Deficiency Drives Chronic Glial Activation and Persistent Astrogliosis in the Retina

At day 30p.i., retinal cross-sections from WT and CD40−/− mice were assessed for glial activation by H&E, Iba1, and GFAP staining (Fig. 11, 12). WT retinas showed substantially resolved inflammation with residual Iba1+ cells adopting a ramified, surveilling morphology and minimal GFAP expression (Fig. 11C, E). In contrast, CD40−/− retinas exhibited significantly elevated Iba1+ cell density with an amoeboid, activated phenotype and markedly increased GFAP expression extending throughout the retinal layers (Fig. 11D, F, G, H). Double immunofluorescence confirmed extensive co-localization of Iba1 and GFAP reactivity in CD40−/− retinas at day 30 p.i. (Fig. 12), indicating a profound and sustained neuroinflammatory environment in the absence of CD40-mediated immune regulation

**Figure 11:**
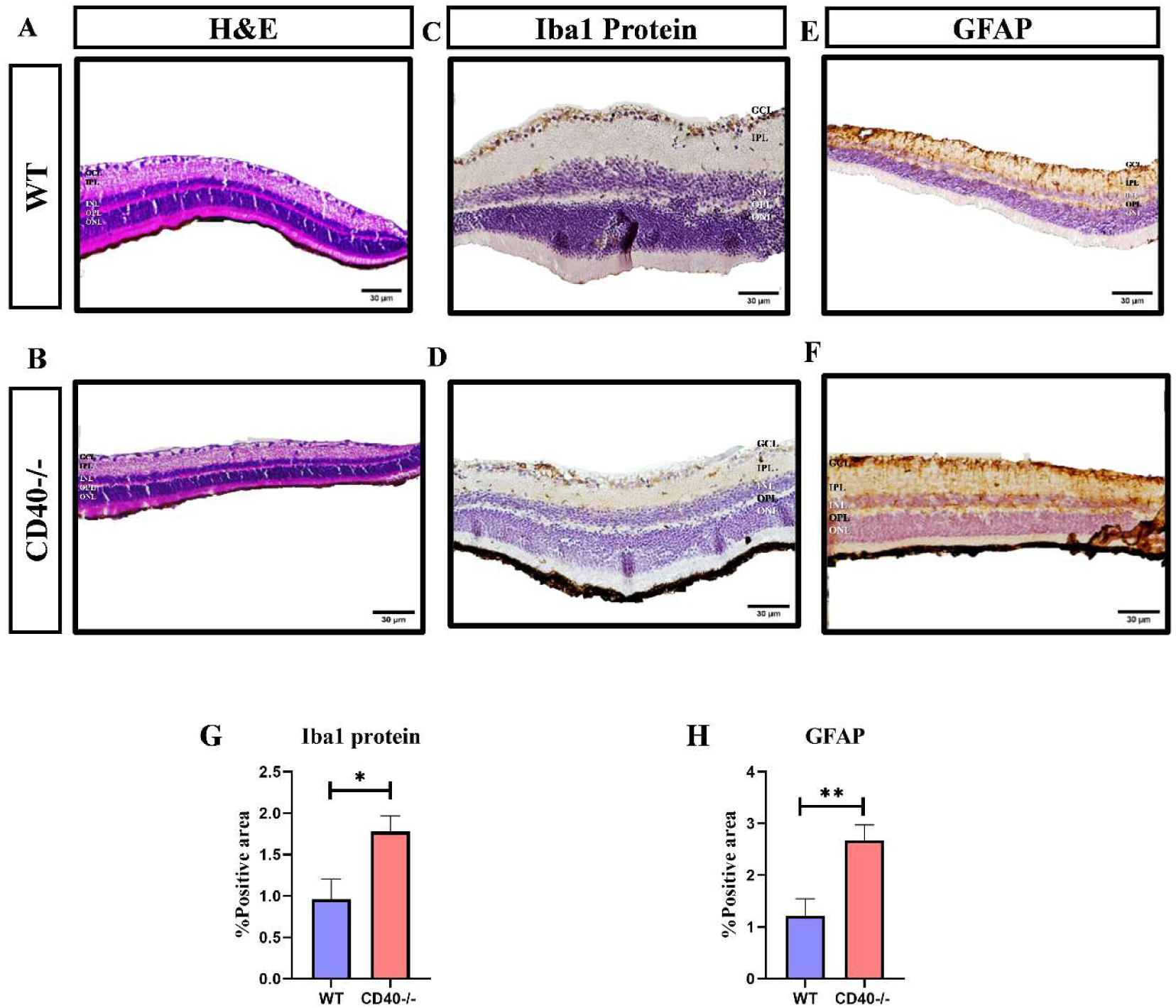
CD40 deficiency drives severe retinal microglial/macrophage activation and persistent astrocyte activation during chronic infection. At day 30 p.i., 5 μm-thick cross-sections of retinas from RSA59-infected (2500 PFU) WT and CD40⁻/⁻ mice were stained with hematoxylin and eosin, and immunohistochemically stained with anti-Iba1 to assess microglial/macrophage activation and anti-GFAP to assess astrogliosis. Representative images show H&E-stained sections (A, B), and a further increase in phagocytic microglia/macrophages in the CD40-/- retina (D) compared to WT (C) was observed. A significant increase in GFAP expression was observed in CD40⁻/⁻ mice (F), extending throughout the retinal sections, compared to WT (E), indicating enhanced astrocyte activation. Quantification of Iba1 and GFAP protein expression is graphically represented in Panels G and H, respectively. The WT and CD40-/- retinal cross sections were magnified at 40X, and the scale bar for all sections is 30 μm. Data are represented as mean ± SEM (N=3 per group). Statistical significance was determined using an unpaired T test; *p < 0.05, **p < 0.01.

**Figure 12:**
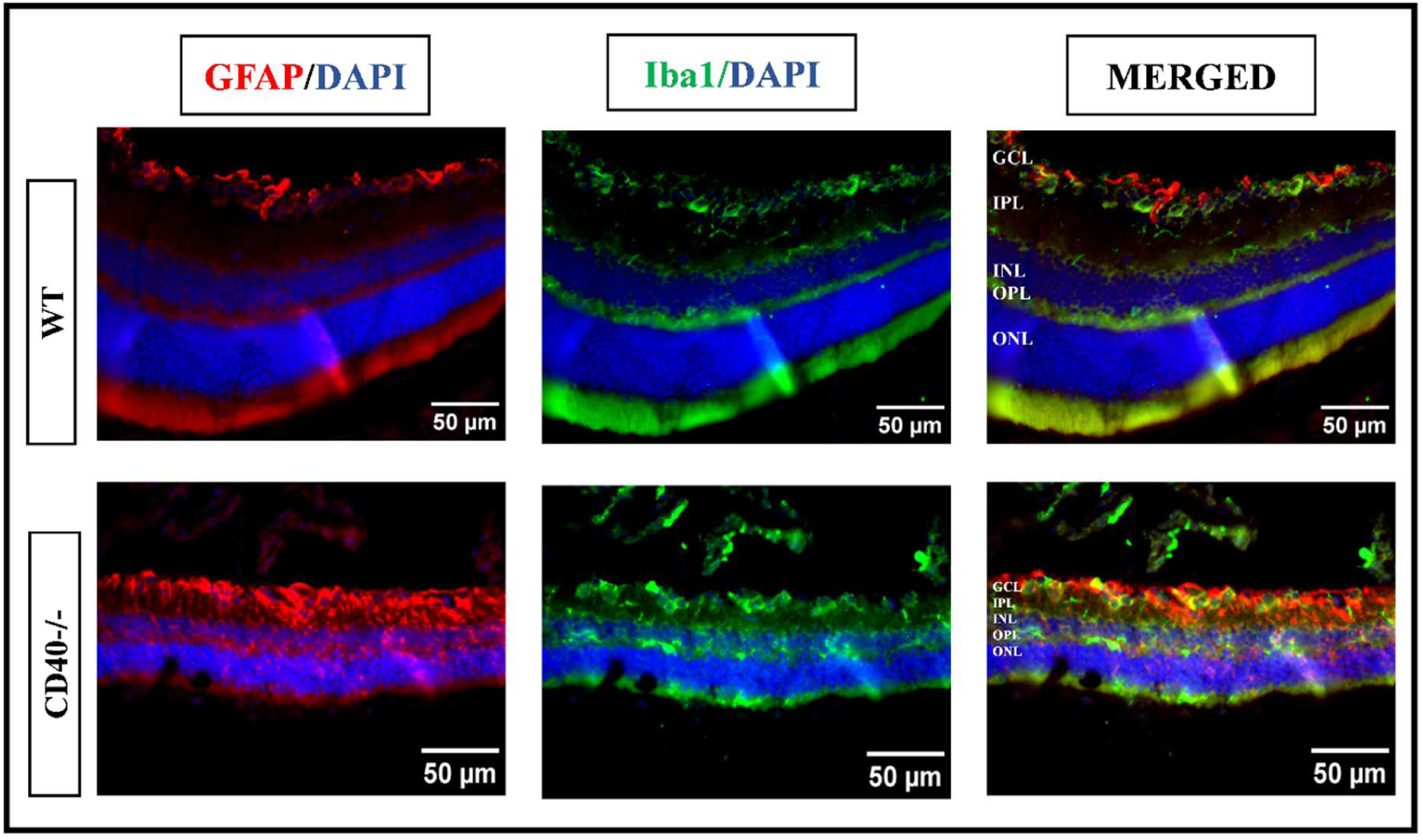
CD40 deficiency leads to significant glial cell activation in the Retina during the chronic stage of infection. At day 30 p.i., cross-sections of RSA59-infected WT and CD40⁻/⁻ mouse retinas were stained to visualize microglia/macrophage activation [Iba1 (green)], astrogliosis [GFAP (red)], and nuclei [DAPI (blue)]. Representative images showed increased glial cell activation in the CD40-/- retina (B) compared with WT (A). Merged images show areas of overlap in staining for GFAP and Iba1. The scale bar for retina sections is 50 µm. Showing one representative image from one of the three mice in one of the three independent experiments.

### CD40 Deficiency Leads to Viral Persistence and Significant Retinal Ganglion Cell Loss in the Chronic Phase

To determine the ultimate neurological consequences of CD40 deficiency in the visual system, retinal sections from day-30p.i. mice were co-stained for viral nucleocapsid protein and the RGC-specific marker Brn3a (Fig. 13). Strikingly, CD40−/− retinas showed significant residual viral N-protein immunoreactivity at day 30p.i., whereas WT retinas had cleared detectable antigen (Fig. 13A, B). Critically, quantification of Brn3a+ neurons in standardized retinal fields revealed a highly significant reduction in the mean fluorescence intensity (MFI) in the RGCs in CD40−/− retinas compared to WT controls (Fig. 13C). This significant reduction in the MFI of Brn3a occurred in the context of persistent viral antigen and severe, unresolved glial activation, indicating that the combined effects of direct viral cytopathic damage and bystander neuroinflammatory injury drive the progressive and irreversible degeneration of retinal neurons in the absence of CD40-mediated innate immune control.

**Figure 13:**
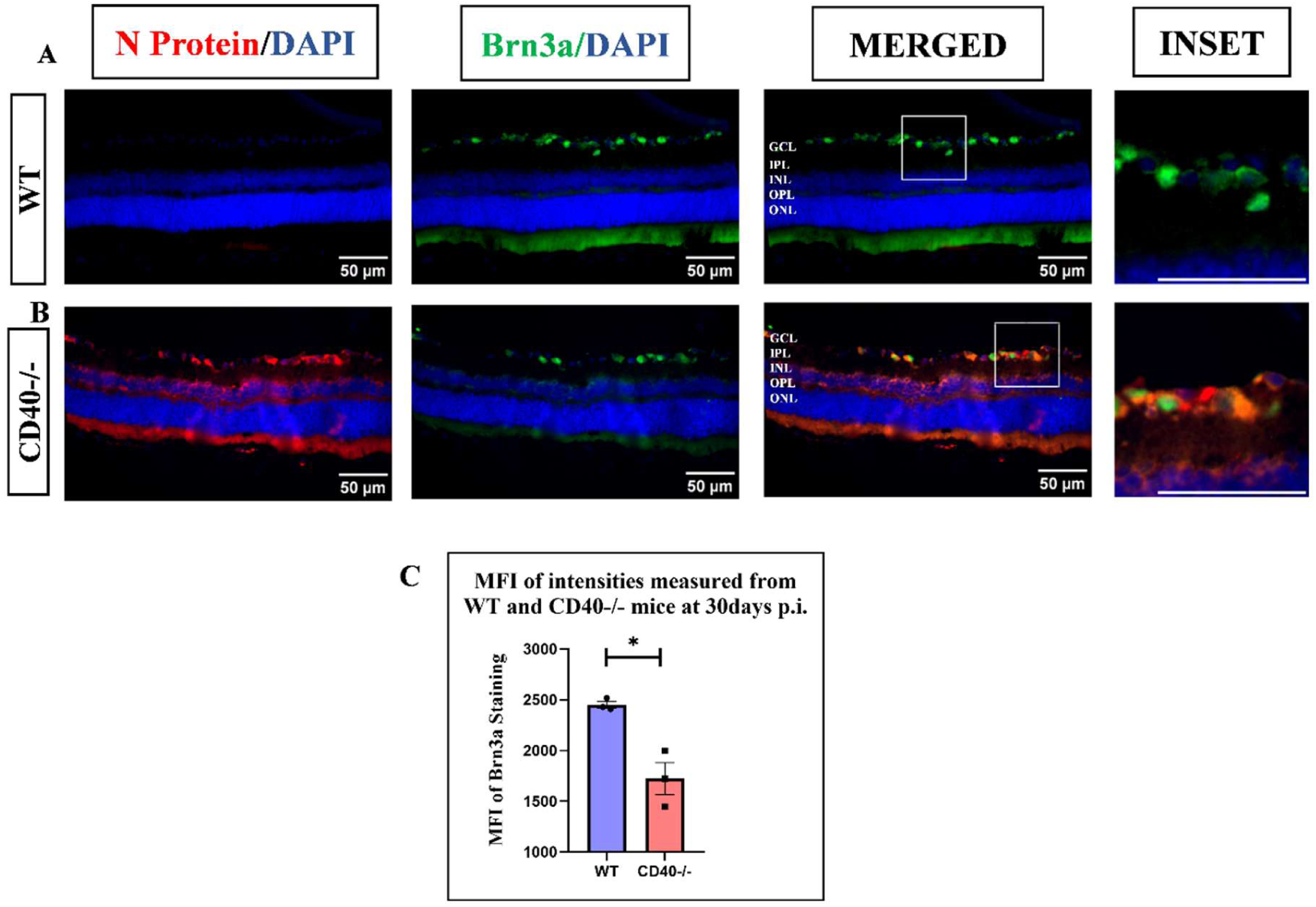
CD40 deficiency leads to viral persistence and RGC loss in the retina in the chronic phase. 5μm longitudinal paraffin sections of optic nerves of day 30 post RSA59-infected WT and CD40^-/-^ mice were stained for viral protein [Nucleocapsid (red)] and RGC marker [Brm3a (green)], and nuclei [DAPI (blue)]. Representative images show increased RGC loss in CD40-/- retina (B), as indicated by Brn3a loss compared with WT (A). Notable viral persistence is observed in CD40-/- retina compared to WT, as indicated by Anti-N staining. Merged images show areas of overlapping staining with Anti-N and Brn3a, respectively. Quantification of Brn3a expression is graphically represented in Panel C. The scale bar for retina sections is 50 µm. Showing one representative image from one of the three mice in one of the three independent experiments. Statistical significance was determined using an unpaired T test; *p < 0.05.

## Discussion

The present study identifies CD40 as a non-redundant component of the innate immune defense of the visual system against neurotropic murine β-coronavirus RSA59. Using the established RSA59 model of viral optic neuritis, we demonstrate that CD40 deficiency triggers a cascade of immunological and pathological events culminating in severe optic nerve demyelination, axonal loss, and a significant decrease in Brn3a MFI, suggesting either cellular stress, metabolic dysfunction, or a commitment to apoptotic cell death(32, 33). These findings extend the neuroimmune framework established by our previous work on the CD40–CD40L-Ifit2 axis to the specialized anatomical context of the optic nerve and retina(22–24).

A critical and novel finding of this study is the characterization of a biphasic glial failure in CD40−/− mice. During the acute phase, CD40−/− mice failed to mount a robust microglia/macrophage response in the optic nerve and retina, despite harboring significantly higher viral loads than WT mice. This counterintuitive dissociation, increased viral burden paired with reduced microglial activation, parallels the phenotype previously observed in Ifit2−/− mice in the brain and spinal cord, where a failure of microglial activation was attributed to dysregulated CX3CR1-dependent innate immune signaling(22, 23). The present data suggest that CD40 plays an analogous role in the visual system, where CD40-mediated signaling in microglia and macrophages is required for early sensing and the mobilization of innate immune cells to sites of viral entry, specifically the RGC layer of the retina. Critically, in the absence of this early microglial response, RSA59 exploits retrograde axonal transport mechanisms far more efficiently, penetrating the GCL barrier and invading the INL, OPL, and ONL layers, a degree of pan-retinal spread not observed in WT mice. This finding is consistent with prior studies demonstrating that RSA59 is normally confined to the GCL in immunocompetent mice and that cell-to-cell fusion mediated by the spike protein fusion peptide is required for deeper retinal penetration(11, 19).

Paradoxically, the same CD40−/− mice that showed impaired microglial activation concurrently exhibited markedly elevated astrogliosis (GFAP+) in both the optic nerve and the retina at the acute and bridging phases. This divergence in the behavior of the two principal glial populations impaired microglial response with compensatory astrocyte activation suggests that reactive astrogliosis may represent a glial substitute response attempting to compensate for the failure of CD40-dependent microglial innate immunity, but that is ultimately insufficient to control viral replication or prevent chronic neurodegeneration(34, 35). The mechanistic basis for this dichotomy may lie in the fact that astrocyte activation does not require CD40 signaling to the same extent as microglia, and that STAT3-dependent astrogliosis can be triggered by a broad range of viral, inflammatory, and damage-associated signals present in the infected CNS, even without intact CD40(36–38). Importantly, the failure of CD40-mediated microglial priming might lead to reduced secretion of pro-inflammatory cytokines (such as TNF-α, IL-12, IL-6), which are normally downstream of CD40 signaling in microglia, depriving the retinal microenvironment of key immunomodulatory signals needed to orchestrate an effective early antiviral response(39–41).

By the chronic stage (day 30p.i.), CD40−/− mice developed a second wave of severe, dysregulated neuroinflammation. The transition from the initial innate failure to a late hyperinflammatory state is reflected in a marked shift in microglia toward an amoeboid phagocytic phenotype, with upregulated expression of P2Y6, TREM2, and CD206. This phagocytic microglial signature has previously been documented in white matter regions of demyelinating spinal cords in the RSA59 model and is consistent with microglia-driven active myelin stripping, a process that amplifies demyelination beyond the direct effects of viral cytopathology(24, 29). In parallel, persistent astrogliosis persisted unabated at day 30 p.i. in CD40−/− retinas and optic nerves, reflecting a chronic, non-resolving neuroinflammatory state driven by the persistent viral RNA reservoir identified by qRT-PCR. The detection of viral RNA without corresponding N-protein immunoreactivity at day 30 is a particularly significant observation that mirrors findings in CD40L−/− mice in our prior studies(24) and suggests that sub-threshold viral replication or the presence of viral genomic fragments maintains continuous innate immune activation even in the apparent absence of detectable antigen.

The present findings on CD40 complement and diverge from our previous characterization of CD40L deficiency in this model in important ways. Both CD40 and CD40L deficiencies impair viral clearance in the acute phase and lead to severe chronic demyelination. In the present study, CD40 deficiency in the visual system specifically impaired microglial activation (Iba1+) while simultaneously driving enhanced astrogliosis, a phenotype distinct from the overt neuroinflammation previously described in the brain. This tissue-specific difference may reflect the unique immunological microenvironment of the eye, where the density and activation threshold of resident microglia differ from those in the brain parenchyma, and where astrocyte populations play a more prominent structural and barrier role. These findings reveal that CD40-mediated immune regulation exhibits anatomical specificity within the CNS, with distinct cellular consequences in the visual system compared with other brain regions.

The loss of Brn3a+ RGC staining in CD40−/− mice at day 30p.i. represents the most clinically impactful finding of this study. RGC loss is the final common pathway linking optic nerve neuroinflammation and demyelination to permanent visual impairment. Our data establish that this neuronal death in CD40−/− mice arises from the convergence of at least three distinct mechanisms: (1) direct viral cytopathic effects, as evidenced by persistent N-protein immunoreactivity in CD40−/− retinas at day 30; (2) bystander neuroinflammatory damage from unresolved microglial and astroglial activation; and (3) retrograde degeneration secondary to severe axonal loss (NFM loss) and demyelination in the optic nerve. This multi-mechanism convergence is consistent with the RGC loss previously reported in RSA59(PP)-infected WT mice, compared with the attenuated loss in fusion-deficient RSA59(P)-infected mice, in which the reduction in retrograde viral transport directly correlated with reduced neurodegeneration(17, 18). The present study extends this concept by demonstrating that a host immune factor, CD40, is equally required to prevent the full cascade of retrograde transport, viral persistence, and neurodegeneration.

An important conceptual advance of this study is the insight it provides into the potentially independent or interconnected roles of CD40–CD40L and the Ifit2 neuroimmune axis in RSA59-induced CNS pathology. Within this circuit, Ifit2 is proposed to function downstream of CD40–CD40L co-stimulation. The similar phenotypic features shared between CD40−/− and Ifit2−/− mice (*Manuscript under preparation*) in the visual system, both showing impaired early microglial activation, enhanced retrograde viral spread, viral persistence, and severe RGC loss, strongly support the hierarchical relationship between these two molecules and demonstrate that both nodes of this axis are independently required in the optic nerve and retina. Disruption at either node of this circuit, CD40 deficiency (as shown here) or Ifit2 deficiency, produces strikingly convergent visual system pathology, further validating the concept that this neuroimmune axis functions as an integrated unit.

The present findings have translational relevance beyond the murine coronavirus model. SARS-CoV-2, the causative agent of COVID-19, is a β-coronavirus sharing structural and functional homologies with MHV/RSA59(42, 43), and neurological sequelae, including optic neuritis and visual disturbances, have been reported in COVID-19 patients and in those with Long COVID(44–46). Mapping the CD40-dependent innate immune pathway that restricts coronavirus retrograde transport and RGC neurodegeneration may provide therapeutic targets relevant to COVID-19-associated neurological disease. Additionally, the MHV model’s capacity to recapitulate both the inflammatory demyelination of MS and the viral etiology hypothesis of MS makes the CD40 pathway identified here relevant to understanding the heterogeneous immune mechanisms in MS-associated optic neuritis (47).

Finally, this study reinforces a growing body of evidence that, in the context of viral CNS infection, the co-stimulatory molecules CD40 and CD40L play protective rather than pathogenic roles, fundamentally distinct from their well-established pathogenic contributions in autoimmune demyelinating disease models such as EAE(24, 48–51). This dichotomy has profound implications for the design of therapeutic strategies targeting CD40 signaling: interventions that may be beneficial in autoimmune neuroinflammation could be detrimental in viral neuroinflammation by impairing the very innate immune circuits required for viral clearance and neuroprotection.

In conclusion, this study demonstrates that CD40, a co-stimulatory receptor of the tumor necrosis factor receptor superfamily expressed on CNS-resident microglia and macrophages, is an essential innate immune regulator that restricts retrograde axonal dissemination of the neurotropic murine β-coronavirus RSA59 in the visual system. CD40 deficiency results in a stereotyped, biphasic pathology; an early innate immune failure characterized by impaired microglial activation concurrent with compensatory astrogliosis, followed by a chronic, dysregulated neuroinflammatory state driven by persistent viral RNA, phagocytic microglia-mediated demyelination, and sustained reactive astrogliosis. This two-phase immune failure culminates in extensive optic nerve demyelination, severe axonal loss, oligodendrocyte depletion/myelin loss, and RGC loss. The CD40-associated neuroimmune axis establishes that the integrity of this circuit is indispensable for neuroprotection in the retina and optic nerve during viral challenge. These findings provide a mechanistic framework for understanding coronavirus-induced optic neuritis and identify CD40 as a potential therapeutic target in infectious optic neuropathies.

## Abbreviations

ON: Optic Neuritis
MS: Multiple Sclerosis
MHV: Mouse Hepatitis Virus
RSA59: Recombinant Spike-gene recombinant of MHV-A59
RGC: Retinal Ganglion Cell
GCL: Ganglion Cell Layer
IPL: Inner Plexiform Layer
INL: Inner Nuclear Layer
OPL: Outer Plexiform Layer
ONL: Outer Nuclear Layer
CNS: Central Nervous System
p.i.: Post-Infection
DM: Demyelinating
NDM: Non-Demyelinating
LGN: Lateral Geniculate Nucleus
WT: Wild-Type
Iba1: Ionized Calcium-Binding Adapter Molecule 1
GFAP: Glial Fibrillary Acidic Protein
MBP: Myelin Basic Protein
PLP: Proteolipid Protein
NFM: Neurofilament Medium Chain
LFB: Luxol Fast Blue
H&E: Hematoxylin and Eosin
qRT-PCR: Quantitative Reverse-Transcription Polymerase Chain Reaction
Brn3a: Brain-Specific Homeobox/POU Domain Protein 3A
ISG: Interferon-Stimulated Gene
Ifit2: Interferon-Induced Protein with Tetratricopeptide Repeats 2
EAE: Experimental Autoimmune Encephalomyelitis
TREM2: Triggering Receptor Expressed on Myeloid Cells-2
PFU: Plaque-Forming Units
EGFP: Enhanced Green Fluorescent Protein
DAB: 3,3′-Diaminobenzidine
PFA: Paraformaldehyde
DAPI: 4′,6-Diamidino-2-Phenylindole

## Declarations

### Ethics Approval and Consent to Participate

All animal procedures were conducted in compliance with institutional ethical guidelines and were approved by the Institutional Animal Ethics Committee (IAEC; IISERK/IAEC/AP/2024/128, Understanding the immunomodulatory role of the immune checkpoint regulator protein CD40 in a murine beta-coronavirus-induced demyelination model). No human subjects were involved in this study. Experiments were performed in accordance with the guidelines of the Committee for Control and Supervision of Experiments on Animals (CCSEA), India.

### Availability of Data and Materials

The data supporting the study findings can be obtained from the corresponding author on request.

### Authors’ Contributions

*Niveditha E*: Data curation, Formal analysis, Investigation, Methodology, Software, Visualization, Writing – original draft, Writing – review & editing

*Bishal Hazra*: Conceptualization, Data curation, Formal analysis, Investigation, Methodology, Software, Visualization, Writing – original draft, Writing – review & editing

*Souvik Karmakar*: Methodology

*Subhajit Das Sarma*: Methodology

*Kenneth S Schindler*: Writing – review & editing

*Jayasri Das Sarma*: Conceptualization, Formal analysis, Funding acquisition, Investigation, Project administration, Resources, Supervision, Validation, Visualization, Writing – original draft, Writing – review & editing

## Acknowledgments

The authors gratefully acknowledge Dr. Julian Leibowitz (Texas A&M University) for providing the anti-nucleocapsid monoclonal antibody (clone 1-16-1) and Judith B. Grinspan (CHOP, Philadelphia) for the PLP and MBP antibodies. We acknowledge the support of the Department of Biological Sciences and the State-of-the-Art Animal Facility at IISER Kolkata, the Intramural and Academic Research Fund of IISER Kolkata. We would also like to acknowledge the University Grants Commission (UGC) and the Council for Scientific and Industrial Research (CSIR) for providing fellowships to BH and SK.

## References

1. Toosy AT, Mason DF, Miller DH. 2014. Optic neuritis. The Lancet. Neurology 13:83–99.

2. Ciapã MA, Șalaru DL, Stătescu C, Sascău RA, Bogdănici CM. 2022. Optic Neuritis in Multiple Sclerosis: A Review of Molecular Mechanisms Involved in the Degenerative Process. Current issues in molecular biology 44:3959–3979.

3. Hoorbakht H, Bagherkashi F. 2012. Optic neuritis, its differential diagnosis and management. The Open Ophthalmology Journal 6:65–72.

4. Shindler KS, Ventura E, Dutt M, Rostami A. 2008. Inflammatory demyelination induces axonal injury and retinal ganglion cell apoptosis in experimental optic neuritis. Experimental Eye Research 87:208–13.

5. Leibowitz U, Alter M. 1968. Optic nerve involvement and diplopia as initial manifestations of multiple sclerosis. Acta neurologica Scandinavica 44:70–80.

6. Constantinescu CS, Farooqi N, O’Brien K, Gran B. 2011. Experimental autoimmune encephalomyelitis (EAE) as a model for multiple sclerosis (MS). British journal of pharmacology 164:1079–106.

7. Sekyi MT, Feri M, Desfor S, Atkinson KC, Golestany B, Beltran F, Tiwari-Woodruff SK. 2024. Demyelination and neurodegeneration early in experimental autoimmune encephalomyelitis contribute to functional deficits in the anterior visual pathway. Scientific Reports 14:24048.

8. Virtanen JO, Jacobson S. 2012. Viruses and multiple sclerosis. CNS & neurological disorders drug targets 11:528–44.

9. Tarlinton RE, Martynova E, Rizvanov AA, Khaiboullina S, Verma S. 2020. Role of Viruses in the Pathogenesis of Multiple Sclerosis. Viruses 12.

10. Shindler KS, Kenyon LC, Dutt M, Hingley ST, Das Sarma J. 2008. Experimental Optic Neuritis Induced by a Demyelinating Strain of Mouse Hepatitis Virus. Journal of Virology 82:8882–8886.

11. Shindler KS, Chatterjee D, Biswas K, Goyal A, Dutt M, Nassrallah M, Khan RS, Das Sarma J. 2011. Macrophage-Mediated Optic Neuritis Induced by Retrograde Axonal Transport of Spike Gene Recombinant Mouse Hepatitis Virus. Journal of Neuropathology & Experimental Neurology 70:470–480.

12. Lavi E, Gilden DH, Wroblewska Z, Rorke LB, Weiss SR. 1984. Experimental demyelination produced by the A59 strain of mouse hepatitis virus. Neurology 34:597–603.

13. Matthews AE, Weiss SR, Paterson Y. 2002. Murine hepatitis virus -- a model for virus-induced CNS demyelination. J Neurovirol 8:76–85.

14. Das Sarma J, Kenyon LC, Hingley ST, Shindler KS. 2009. Mechanisms of Primary Axonal Damage in a Viral Model of Multiple Sclerosis. The Journal of Neuroscience : the official journal of the Society for Neuroscience 29:10272–10280.

15. Das Sarma J, Fu L, Tsai JC, Weiss SR, Lavi E. 2000. Demyelination determinants map to the spike glycoprotein gene of coronavirus mouse hepatitis virus. J Virol 74:9206–13.

16. Das Sarma J, Iacono K, Gard L, Marek R, Kenyon LC, Koval M, Weiss SR. 2008. Demyelinating and nondemyelinating strains of mouse hepatitis virus differ in their neural cell tropism. J Virol 82:5519–26.

17. Singh M, Kishore A, Maity D, Sunanda P, Krishnarjuna B, Vappala S, Raghothama S, Kenyon LC, Pal D, Das Sarma J. 2019. A proline insertion-deletion in the spike glycoprotein fusion peptide of mouse hepatitis virus strongly alters neuropathology. Journal of Biological Chemistry 294:8064–8087.

18. Rout SS, Singh M, Shindler KS, Das Sarma J. 2020. One proline deletion in the fusion peptide of neurotropic mouse hepatitis virus (MHV) restricts retrograde axonal transport and neurodegeneration. J Biol Chem 295:6926–6935.

19. Singh M, Khan RS, Dine K, Das Sarma J, Shindler KS. 2018. Intracranial Inoculation Is More Potent Than Intranasal Inoculation for Inducing Optic Neuritis in the Mouse Hepatitis Virus-Induced Model of Multiple Sclerosis. Front Cell Infect Microbiol 8:311.

20. Chatterjee D, Addya S, Khan RS, Kenyon LC, Choe A, Cohrs RJ, Shindler KS, Sarma JD. 2014. Mouse Hepatitis Virus Infection Upregulates Genes Involved in Innate Immune Responses. PLOS ONE 9:e111351.

21. Syage A, Pachow C, Cheng Y, Mangale V, Green KN, Lane TE. 2023. Microglia influence immune responses and restrict neurologic disease in response to central nervous system infection by a neurotropic murine coronavirus. Front Cell Neurosci 17:1291255.

22. Das Sarma J, Burrows A, Rayman P, Hwang MH, Kundu S, Sharma N, Bergmann C, Sen GC. 2020. Ifit2 deficiency restricts microglial activation and leukocyte migration following murine coronavirus (m-CoV) CNS infection. PLoS Pathog 16:e1009034.

23. Sharma M, Chakravarty D, Hussain A, Zalavadia A, Burrows A, Rayman P, Sharma N, Kenyon LC, Bergmann C, Sen GC, Das Sarma J. 2023. Ifit2 restricts murine coronavirus spread to the spinal cord white matter and its associated myelin pathology. J Virol 97:e0074923.

24. Saadi F, Chakravarty D, Kumar S, Kamble M, Saha B, Shindler KS, Das Sarma J. 2021. CD40L protects against mouse hepatitis virus-induced neuroinflammatory demyelination. PLoS Pathog 17:e1010059.

25. Das Sarma J, Scheen E, Seo SH, Koval M, Weiss SR. 2002. Enhanced green fluorescent protein expression may be used to monitor murine coronavirus spread in vitro and in the mouse central nervous system. J Neurovirol 8:381–91.

26. McGavern DB, Murray PD, Rodriguez M. 1999. Quantitation of spinal cord demyelination, remyelination, atrophy, and axonal loss in a model of progressive neurologic injury. Journal of Neuroscience Research 58:492–504.

27. Wheeler DL, Sariol A, Meyerholz DK, Perlman S. 2018. Microglia are required for protection against lethal coronavirus encephalitis in mice. J Clin Invest 128:931–943.

28. Donnelly DJ, Gensel JC, Ankeny DP, van Rooijen N, Popovich PG. 2009. An efficient and reproducible method for quantifying macrophages in different experimental models of central nervous system pathology. Journal of Neuroscience Methods 181:36–44.

29. Kamble M, Saadi F, Kumar S, Saha B, Das Sarma J. 2023. Inducible nitric oxide synthase deficiency promotes murine-β-coronavirus induced demyelination. Virology Journal 20:51.

30. Santos AM, Calvente R, Tassi M, Carrasco M-C, Martín-Oliva D, Marín-Teva JL, Navascués J, Cuadros MA. 2008. Embryonic and postnatal development of microglial cells in the mouse retina. The Journal of Comparative Neurology 506:224–239.

31. Chakravarty D, Saadi F, Kundu S, Bose A, Khan R, Dine K, Kenyon LC, Shindler KS, Das Sarma J. 2020. CD4 Deficiency Causes Poliomyelitis and Axonal Blebbing in Murine Coronavirus-Induced Neuroinflammation. J Virol 94.

32. Nadal-Nicolás FM, Jiménez-López M, Sobrado-Calvo P, Nieto-López L, Cánovas-Martínez I, Salinas-Navarro M, Vidal-Sanz M, Agudo M. 2009. Brn3a as a Marker of Retinal Ganglion Cells: Qualitative and Quantitative Time Course Studies in Naïve and Optic Nerve–Injured Retinas. Investigative Ophthalmology & Visual Science 50:3860–3868.

33. Meng M, Chaqour B, O’Neill N, Dine K, Sarabu N, Ying G-S, Shindler K, Ross A. 2024. Comparison of Brn3a and RBPMS Labeling to Assess Retinal Ganglion Cell Loss During Aging and in a Model of Optic Neuropathy. Investigative Ophthalmology & Visual Science 65:19.

34. Sofroniew MV, Vinters HV. 2010. Astrocytes: biology and pathology. Acta Neuropathol 119:7–35.

35. Liddelow SA, Barres BA. 2017. Reactive Astrocytes: Production, Function, and Therapeutic Potential. Immunity 46:957–967.

36. Herrmann JE, Imura T, Song B, Qi J, Ao Y, Nguyen TK, Korsak RA, Takeda K, Akira S, Sofroniew MV. 2008. STAT3 is a critical regulator of astrogliosis and scar formation after spinal cord injury. J Neurosci 28:7231–43.

37. Acarin L, Gonzແlez B, Castellano B. 2000. STAT3 and NFκB Activation Precedes Glial Reactivity in the Excitotoxically Injured Young Cortex but not in the Corresponding Distal Thalamic Nuclei. Journal of Neuropathology & Experimental Neurology 59:151–163.

38. Hamby ME, Sofroniew MV. 2010. Reactive astrocytes as therapeutic targets for CNS disorders. Neurotherapeutics 7:494–506.

39. Aloisi F, Penna G, Polazzi E, Minghetti L, Adorini L. 1999. CD40-CD154 interaction and IFN-gamma are required for IL-12 but not prostaglandin E2 secretion by microglia during antigen presentation to Th1 cells. J Immunol 162:1384–91.

40. Tan J, Town T, Paris D, Mori T, Suo Z, Crawford F, Mattson MP, Flavell RA, Mullan M. 1999. Microglial Activation Resulting from CD40-CD40L Interaction After β-Amyloid Stimulation. Science (New York, N.Y.) 286:2352–2355.

41. Gerritse K, Laman JD, Noelle RJ, Aruffo A, Ledbetter JA, Boersma WJ, Claassen E. 1996. CD40-CD40 ligand interactions in experimental allergic encephalomyelitis and multiple sclerosis. Proc Natl Acad Sci U S A 93:2499–504.

42. Grabherr S, Ludewig B, Pikor NB. 2021. Insights into coronavirus immunity taught by the murine coronavirus. Eur J Immunol 51:1062–1070.

43. Safiriyu AA, Mulchandani V, Anakkacheri MN, Pal D, Das Sarma J. 2023. Proline-Proline Dyad in the Fusion Peptide of the Murine β-Coronavirus Spike Protein’s S2 Domain Modulates Its Neuroglial Tropism. Viruses 15.

44. Georganta I, Chasapi D, Smith CJ, Kopsidas K, Tatham A. 2023. Systematic review exploring the clinical features of optic neuritis after SARS-CoV infection and vaccination. BMJ Open Ophthalmol 8.

45. Shi J, Danesh-Meyer HV. 2024. A review of neuro-ophthalmic sequelae following COVID-19 infection and vaccination. Front Cell Infect Microbiol 14:1345683.

46. Dağ Şeker E, Erbahçeci Timur İ E. 2023. Assessment of early and long-COVID related retinal neurodegeneration with optical coherence tomography. Int Ophthalmol 43:2073–2081.

47. Kishore A, Kanaujia A, Nag S, Rostami AM, Kenyon LC, Shindler KS, Das Sarma J. 2013. Different mechanisms of inflammation induced in virus and autoimmune-mediated models of multiple sclerosis in C57BL6 mice. Biomed Res Int 2013:589048.

48. Aarts S, Seijkens TTP, van Dorst KJF, Dijkstra CD, Kooij G, Lutgens E. 2017. The CD40-CD40L Dyad in Experimental Autoimmune Encephalomyelitis and Multiple Sclerosis. Front Immunol 8:1791.

49. Lu Y, Xu M, Dorrier CE, Zhang R, Mayer CT, Wagner D, McGavern DB, Hodes RJ. 2022. CD40 Drives Central Nervous System Autoimmune Disease by Inducing Complementary Effector Programs via B Cells and Dendritic Cells. The Journal of Immunology 209:2083–2092.

50. Hazra B, Das Sarma J. 2024. The CD40/CD40 ligand dyad and its downstream effector molecule ISG54 in relating acute neuroinflammation with persistent, progressive demyelination. IUBMB Life 76:313–331.

51. Lu Y, Xu M, Dorrier CE, Zhang R, Mayer CT, Wagner D, McGavern DB, Hodes RJ. 2022. CD40 Drives Central Nervous System Autoimmune Disease by Inducing Complementary Effector Programs via B Cells and Dendritic Cells. J Immunol 209:2083–2092.

